# Preproteins couple the intrinsic dynamics of SecA to its ATPase cycle to translocate via a catch and release mechanism

**DOI:** 10.1101/2021.08.31.458331

**Authors:** Srinath Krishnamurthy, Marios-Frantzeskos Sardis, Nikolaos Eleftheriadis, Katerina E. Chatzi, Jochem H. Smit, Konstantina Karathanou, Giorgos Gouridis, Athina G. Portaliou, Ana-Nicoleta Bondar, Spyridoula Karamanou, Anastassios Economou

## Abstract

Protein machines undergo conformational motions to interact with and manipulate polymeric substrates. The Sec translocase promiscuously recognizes, becomes activated and secretes >500 non-folded preprotein clients across bacterial cytoplasmic membranes. Here, we reveal that the intrinsic dynamics of the translocase ATPase, SecA, and of preproteins combine to achieve translocation. SecA possesses an intrinsically dynamic preprotein clamp attached to an equally dynamic ATPase motor. Alternating motor conformations are finely controlled by the γ-phosphate of ATP, while ADP causes motor stalling, independently of clamp motions. Functional preproteins physically bridge these independent dynamics. Their signal peptide promotes clamp closing; their mature domain overcomes the rate limiting ADP release. While repeated ATP cycles shift the motor between unique states, multiple conformationally frustrated prongs in the clamp repeatedly ‘catch and release’ trapped preprotein segments until translocation completion. This universal mechanism allows any preprotein to promiscuously recognize the translocase, usurp its intrinsic dynamics and become secreted.

## Introduction

Protein machines chemically modify, reshape, disaggregate and transport nucleic acids and polypeptides (Avellaneda et al., 2017; Flechsig and Mikhailov, 2019; Kurakin, 2006). In doing so, they convert between auto-inhibited and active states that commonly depend on intrinsic structural dynamics (Nussinov et al., 2018). A fascinating paradigm of such a machine is the bacterial Sec translocase, involved in secretion of client proteins (preproteins) across the inner membrane. Its SecA ATPase subunit, a four domain Superfamily 2 DEAD box helicase, interacts with non-folded signal peptide-bearing clients, nucleotides, lipids, chaperones and the trimeric SecYEG channel (De Geyter et al., 2020; Rapoport et al., 2017; Tsirigotaki et al., 2017a). The coordination of these sub-reactions achieves translocase activation by exploiting a multi-tiered intrinsic dynamics nexus (Corey et al., 2019; Gouridis et al., 2013; Krishnamurthy et al., 2021; Sardis and Economou, 2010). The latter is built on an extensive Hydrogen-bonded (H-bond) network and requires minor energetic input from ligands (Krishnamurthy et al., 2021). While partner subunits and nucleotides regulate and prime the dynamics landscape of the Sec translocase, activation is ultimately only driven by the secretory clients. Loosely conserved client sequence features allow the translocase to bind, get activated by and ultimately translocate ∼500 different clients across the bacterial inner membrane, at the expense of energy (Tsirigotaki et al., 2017a). This occurs via a poorly understood universal activation mechanism that is preprotein sequence-agnostic.

Protein intrinsic dynamics are multi-leveled (Henzler-Wildman et al., 2007; Yang et al., 2014): quaternary motions of subunits, tertiary motions within a single chain, rigid body motions of large structural domains and local motions in small numbers of residues (Krishnamurthy et al., 2021). How intrinsic dynamics couple allostery to function remains unclear (Bhabha et al., 2015; Loutchko and Flechsig, 2020; Zhang et al., 2019) and characterizing it mechanistically is all the more challenging in multi-liganded, multi-partner enzymes such as the Sec translocase, that operate in an apparent hierarchical manner.

Cytoplasmic SecA is dimeric, ADP-bound and quiescent (SecA_2_; Fig. 1A.I) and chaperones preprotein clients (Sianidis et al., 2001). Its helicase motor (comprising <u>N</u>ucleotide <u>B</u>inding <u>D</u>omains 1/2) is fused to an ATPase suppressing C-domain and a <u>P</u>reprotein <u>B</u>inding <u>D</u>omain (PBD) (Fig. S1A) that is rooted via a Stem in NBD1. The PBD intrinsically rotates from a distal inactive “Wide-Open” position towards NBD2 (“Closed” positions) in a crab-claw motion (Ernst et al., 2018; Krishnamurthy et al., 2021; Sardis and Economou, 2010; Vandenberk et al., 2019), to clamp mature domains (Bauer and Rapoport, 2009). Binding to the SecYEG channel (Fig. 1A.II, “primed”) enhances the local dynamics of SecA primarily in the helicase motor and attached Scaffold and Stem [Fig. S1A; (Krishnamurthy et al., 2021)]. Asymmetric binding of SecA_2_ to the channel increases clamp dynamics and interconversion between open and closed states in the channel-bound protomer referred to as “active” (Krishnamurthy et al., 2021). This “primed” translocase has 10-fold higher affinity for preproteins (Gouridis et al., 2009; Gouridis et al., 2013; Hartl et al., 1990) yet, does not substantially turn ATP over (Fak et al., 2004; Keramisanou et al., 2006; Krishnamurthy et al., 2021; Sianidis et al., 2001). The translocase becomes fully activated only after non-folded preproteins bind to it as bivalent ligands with discrete binding sites on SecA for their signal peptides (in the PBD bulb) and mature domains (on and around the PBD Stem) [Fig. S1B; (Chatzi et al., 2017; Gelis et al., 2007; Sardis et al., 2017)]. Signal peptides promote a low activation energy conformation (Fig. 1A.III, “triggered”) that fully activates the ATPase activity of the translocase, presumably overcoming the stable ADP state (Fig. 1A.IV, “activated”), and leading to segmental translocation (Fig. 1A.V, “processive translocation”).

**Figure 1:**
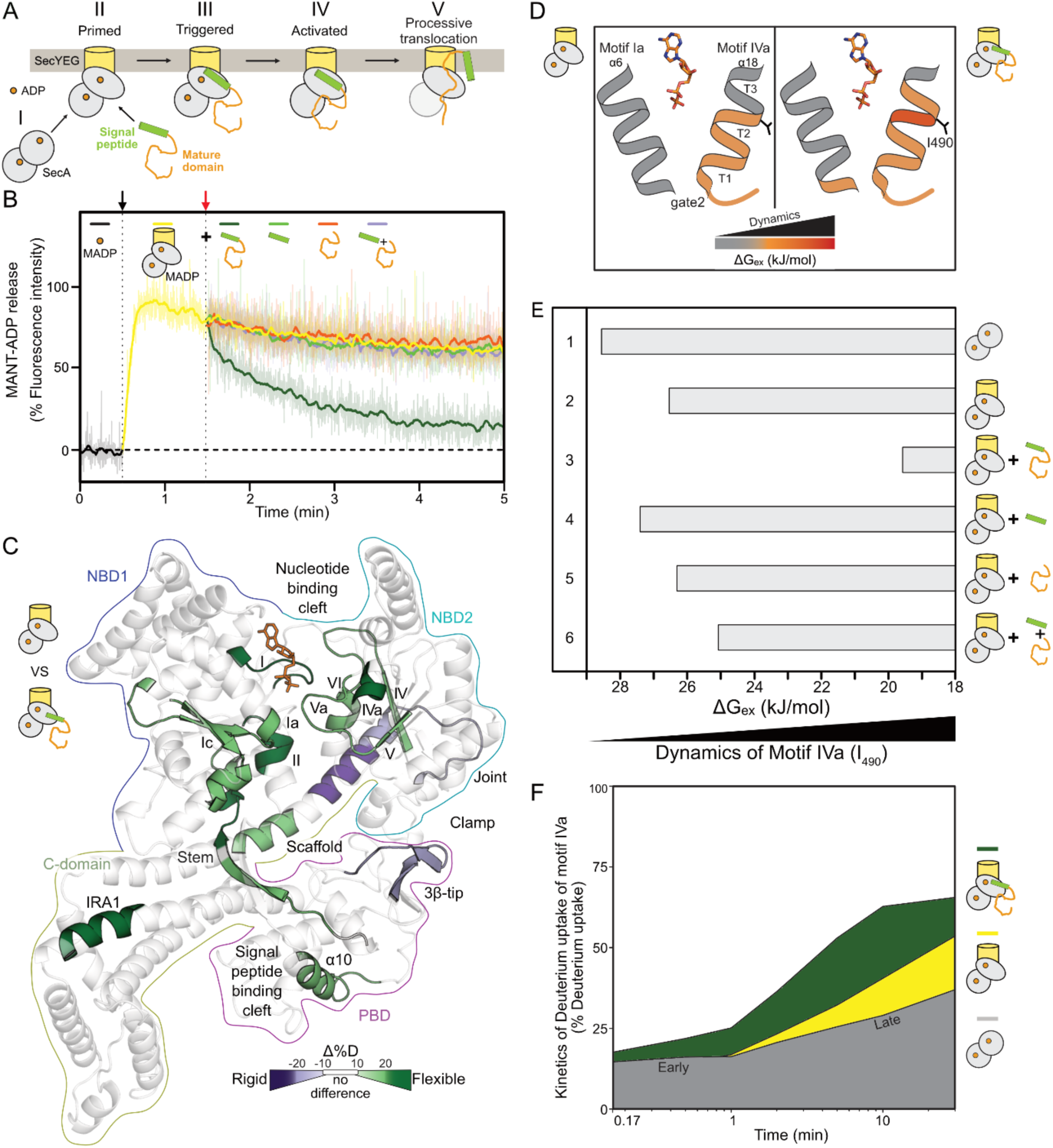
Preproteins induce ADP release by allosterically enhancing the dynamics of the ATPase motor. **A.** Cytoplasmic SecA is an ADP-bound quiescent dimer (I), that binds asymmetrically to the SecYEG channel through the active protomer (II, grey oval). The preprotein is targeted to the translocase through binding of signal peptide and mature domain that act together as a bivalent ligand. Signal peptide binding on the translocase leads to triggering (III). Preprotein binding activates the translocase (IV) for processive ATP hydrolysis cycles that result in preprotein translocation (V). **B.** ADP release assay. The fluorescence intensity of MANT-ADP increases upon binding to channel-primed translocase (1 µM; black arrow at 30 s). The reaction was chased (red arrow; at 90 s) with the indicated ligand. Data were recorded for 5 min. The drop in fluorescence intensity corresponds to the release of MANT-ADP from the nucleotide binding pocket of SecA. Ligands were added at saturating concentrations (see methods). Data are independently normalized using the fluorescence intensity of free MANT-ADP (at 30 s) as 0 %, and the intensity of translocase-bound MANT-ADP (at 45 s) as 100 %. Raw fluorescence traces (transparent lines; *n=3-4*) are presented along with smoothened data (solid lines, LOWESS smoothening). Preprotein: proPhoA_1-122_, mature domain: PhoA_23-122_ (Chatzi et al., 2017). **C.** Effect of proPhoA_1-122_ binding on the local dynamics of SecA_2_:ADP:SecYEG, by HDX-MS (only one protomer shown for simplicity). D-uptake differences between SecA_2_:ADP:SecYEG (top pictogram: “reference”) and SecA_2_:ADP:SecYEG:preprotein (bottom pictogram: “test”) are mapped onto the closed clamp structure of SecA derived from MD simulations with the helicase motor and its gate2 (motifs Ia and IVa) in the open state for better visualization (Krishnamurthy et al., 2021). Decreased/increased dynamics: purple/green respectively; no difference:transparent grey. Domain contours are coloured, ADP: orange sticks. **D.** ΔG_ex_ values representing protein dynamics (covering a range of ) were calculated by PyHDX from HDX-MS data (Smit et al., 2021a) and were mapped onto a cartoon of the gate2 in its closed state with its two helices comprising helicase motifs Ia (α6) and IVa (α18). Dynamics of SecA_2_:ADP:SecYEG in preprotein free (left) and bound (right) state are shown. α18 consists of 3 turns. I_490_ (located between turns 2 and 3) reports on motif IVa dynamics. ADP: orange sticks. See also Fig. S1D. **E.** ΔG_ex_ values (from D) for residue I_490_ (motif IVa) were determined under the indicated conditions. Decreased ΔG_ex_ values correspond to increased dynamics of I_490_. **F.** D-uptake kinetic plots of a motif IVa peptide (aa488-501), shown as a percentage of the full deuteration control (Table S1), for SecA_2_:ADP (grey), SecA_2_:ADP:SecYEG [yellow; (Krishnamurthy et al., 2021)] and SecA_2_:ADP:SecYEG:preprotein (green) are shown. Quiescent SecA_2_:ADP showed biphasic D-uptake kinetics, with a slow initial phase and a fast second one. Data points refer to labeling

We previously developed and use here an integrated, multi-pronged approach to determine the intrinsic dynamics of SecA and probed how these underlie the conversion from a quiescent to a primed SecY-bound state (Karathanou and Bondar, 2019; Krishnamurthy et al., 2021; Vandenberk et al., 2019). We probed ‘global’ dynamics/H-bond networks with atomistic molecular dynamics (MD) simulations and graph analysis, PBD clamp motions by single molecule Förster resonance energy transfer (smFRET) and ‘local’ dynamics by hydrogen deuterium exchange mass spectrometry (HDX-MS). smFRET and HDX-MS experiments are carried out with inverted membrane vesicles, in translocation-permitting conditions identical to the ones used for biochemical dissection (Krishnamurthy et al., 2021) and not in the presence of detergents to avoid monomerizing SecA and altering translocase dynamics (Ahdash et al., 2019; Or et al., 2002).

We now reveal that preproteins achieve their translocation by physically bridging the otherwise largely uncoupled intrinsic dynamics of the SecA helicase motor and those of the preprotein clamp. Signal peptide and mature domain binding on their respective binding sites on the primed translocase [Fig. S1B; (Sardis et al., 2017)] drive specific events: the signal peptide promotes closing of the clamp. This results in enhanced dynamics in the ATPase motor and at the mature domain binding site around the Stem. These local changes facilitate mature domain-driven ADP release. A fresh nucleotide cycle initiates as ATP binds the *apo* motor. Each stage of the hydrolysis cycle leads to distinct states of the ATPase motor that rely on sensing of the *γ*-phosphate. Importantly, these cyclic changes also affect four frustrated prongs that line the preprotein clamp. Dynamics changes lead to specific transient binding on mutliple islands along the client chain resulting in ‘catch and release’ cycles that coincide with active translocation. Thus, through multi-site binding, the elongated client acts as an external temporary physical bridge that couples clamp motions to intrinsic ATPase motor states that are directly regulated by the ATPase cycle. Upon completion of translocation, the client-less machine can no longer overcome the ADP state and becomes quiescent.

## Results

### Preprotein-stimulated ADP release from the helicase motor

Preproteins stimulate ATP turnovers at the translocase. To do so, they must first de-stabilize the robust SecA:ADP state. To test this we used fluorescent MANT-ADP and monitored its binding to/release from the SecYEG:SecA_2_ translocase, in the presence or absence of preprotein (Fig. 1B). The fluorescence intensity of MANT-ADP increases upon binding to the helicase motor of SecA at 37°C (Fig 1B; x axis, black arrow) (Galletto et al., 2005; Karamanou et al., 2005; Krishnamurthy et al., 2021) and remains high through the time course of the experiment (Fig. 1B, yellow line) indicating tight ADP binding. Upon addition of preprotein [proPhoA_1-122_; (Chatzi et al., 2017)], at the indicated time point (Fig. 1B, x axis, orange arrow), the fluorescence intensity drops (green line), indicative of MANT-ADP release.

ADP could not be released from SecA_2_ in solution (Fig. S1C, top) nor at non-physiological temperature (10°C; bottom). Signal peptide (Fig. 1B, green) or mature domain (orange) added alone, or combined *in trans* (purple), at concentrations several times over their K_d_, failed to cause measurable ADP release. Therefore, this reaction is on pathway since only functional preproteins induce ADP release to activate translocase only at physiological conditions.

### Preprotein-enhanced helicase motor dynamics underlie ADP release

Next, we studied whether preprotein-stimulated ADP release and initiation of the translocation reaction correlate with changes in the local intrinsic dynamics of SecYEG:SecA_2_:ADP. For this we used HDX-MS and calculated the per residue Gibbs free energy of exchange (ΔG_ex_) (Krishnamurthy et al., 2021; Smit et al., 2021a). ΔG_ex_ values correlate well with protein intrinsic dynamics and map at residue level flexible (Fig. S1D; low ΔG values; red/orange) or rigid (high ΔG values; transparent grey) protein regions. To quantify the effect of preprotein binding on the dynamics of the translocase we compared the D-uptake between the SecYEG:SecA_2_:ADP:preprotein (Fig. S1D, right) to SecYEG:SecA_2_:ADP as a reference state [left; (Krishnamurthy et al., 2021)]. The resulting ΔD-uptake structural map highlighted that preprotein binding has two main effects. It significantly increased dynamics (Fig. 1C; green hues) at the motor [at helicase motifs (roman numerals) and parallel β-sheets; (Fig. S1E)], the mature domain binding site (IRA1 and Stem) and near the signal peptide binding cleft (PBD_α10_; S1A and D) of SecA. (Keramisanou et al., 2006). In parallel, it decreased dynamics (purple hues) in the Joint/beginning of the Scaffold and the 3β-tip_PBD_. As the effects seen within the helicase motor occur far from the identified preprotein binding sites (Fig. S1B), they are likely allosteric. They are also physiological responses as preprotein binds (Gouridis 2009) but did not significantly alter SecA_2_ dynamics in solution (Table S1).

In the nucleotide cleft motifs I, II and Va_R509_ directly bind the phosphate groups of ATP [(Papanikolau et al., 2007); Fig. S1E, left] and motifs IV and V are important parallel β-strands of the NBD2 core. At the nucleotide cleft periphery, motifs Ia and IVa form the lateral gate2 with defined closed/open states affecting NBD1 and 2 association (Fig. S1E, right)(Papanikolau et al., 2007) and likely regulate access to the nucleotide cleft (see below). Gate2 together with gate1 at the bottom of the nucleotide cleft (Fig. S1E, left), control the onset of ATP hydrolysis (Karamanou et al., 2007). Increased dynamics at these motifs proves that upon preprotein binding, nucleotide contacts in the helicase motor are weakened, presumably underlying ADP release (Fig. 1B).

In conclusion, binding of functional preprotein to the PBD (Chatzi et al., 2017; Gelis et al., 2007) causes long range allosteric effects that affect the dynamics of the motor at several helicase motifs internally and at the peripheral gate2 (Fig. 1D-F), drives ADP release (Fig. 1B) and thus allows initiation of the ATPase cycle.

### Motif IVa of gate2 senses ligands through its intrinsic dynamics

Gate2 is particularly intriguing. Motif IVa has elevated basal dynamics and is sensitive to multiple interactants (Krishnamurthy et al., 2021), is located between the ADP-binding motif Va and the motifs IV and Vb on β-strands of the NBD2 core (Fig. S1E). It comprises a flexible linker followed by a 3-turn helix with a constantly dynamic first half (Fig. 1D, left, orange; S1D, left) followed by a conditionally dynamic second half (turns 2-3). Preprotein binding increased dynamics specifically at turns 2 and 3, around residue I_490_ (Fig. 1D and S1D, right) and this provided us with a powerful assay to quantify dynamics changes in motif IVa (Fig. 1E). Channel binding marginally increases its dynamics compared to the SecA_2_:ADP state (i.e. ΔG_ex_ decreases; Fig. 1E, compare lane 2 to 1; ΔΔG_ex_ = ∼2 kJ/mol), while preprotein binding to the holo-translocase significantly enhances them (lane 3; ΔΔG_ex_ = ∼6 kJ/mol). Addition of signal peptide (lane 4) or mature domain (lane 5) alone, or together *in trans* (lane 6), failed to recapitulate the preprotein effect on motif IVa flexibility.

Motif IVa shows complex dynamics taking up D with biphasic kinetics (Fig. 1F, grey), suggestive of distinct modulation of its energy landscape in two differentially flexible conformational steps. Channel binding selectively increased only the second ‘late’ phase dynamics (yellow), while preprotein added on top, increased both phases (green), suggesting a major reorganization of the conformational landscape of motif IVa. Neither step is sufficient alone; both are necessary for complete activation of the translocase and preprotein could be replaced neither by signal peptide (Fig. S1F.I) nor by mature domain (II) added alone, or together *in trans* (III).

We concluded that motif IVa of gate2 uses its intrinsic dynamics to sense ligands.

### ADP-antagonized, signal peptide-regulated Motif IVa dynamics

We next sought to discern the contribution of each preprotein moiety to activating the transloase, first probing the effect of the signal peptide. The proPhoA signal peptide had marginal effects on the dynamics of the helicase motor of SecYEG:SecA_2_:ADP (Fig. 1E; S1F.I; Table S1). We presumed this to be due to bound ADP antagonizing subtle dynamics effects and tested the effect of signal peptide binding on the *apo* SecYEG:SecA_2_. This time we observed a remarkable increase in extremely localized dynamics at turn 2 of motif IVa and the middle of the Scaffold (Fig. 2A). Time-dependent dynamics of Motif IVa revealed that signal peptide binding increased the dynamics of the early but not the late (channel-dependent) phase (Fig. 2B, compare line to shaded yellow) and the effect was less than that of the preprotein (compare line to shaded green; Fig. S2A). This was corroborated by quantification monitoring of I_490_ (Fig. S2B).

**Figure 2:**
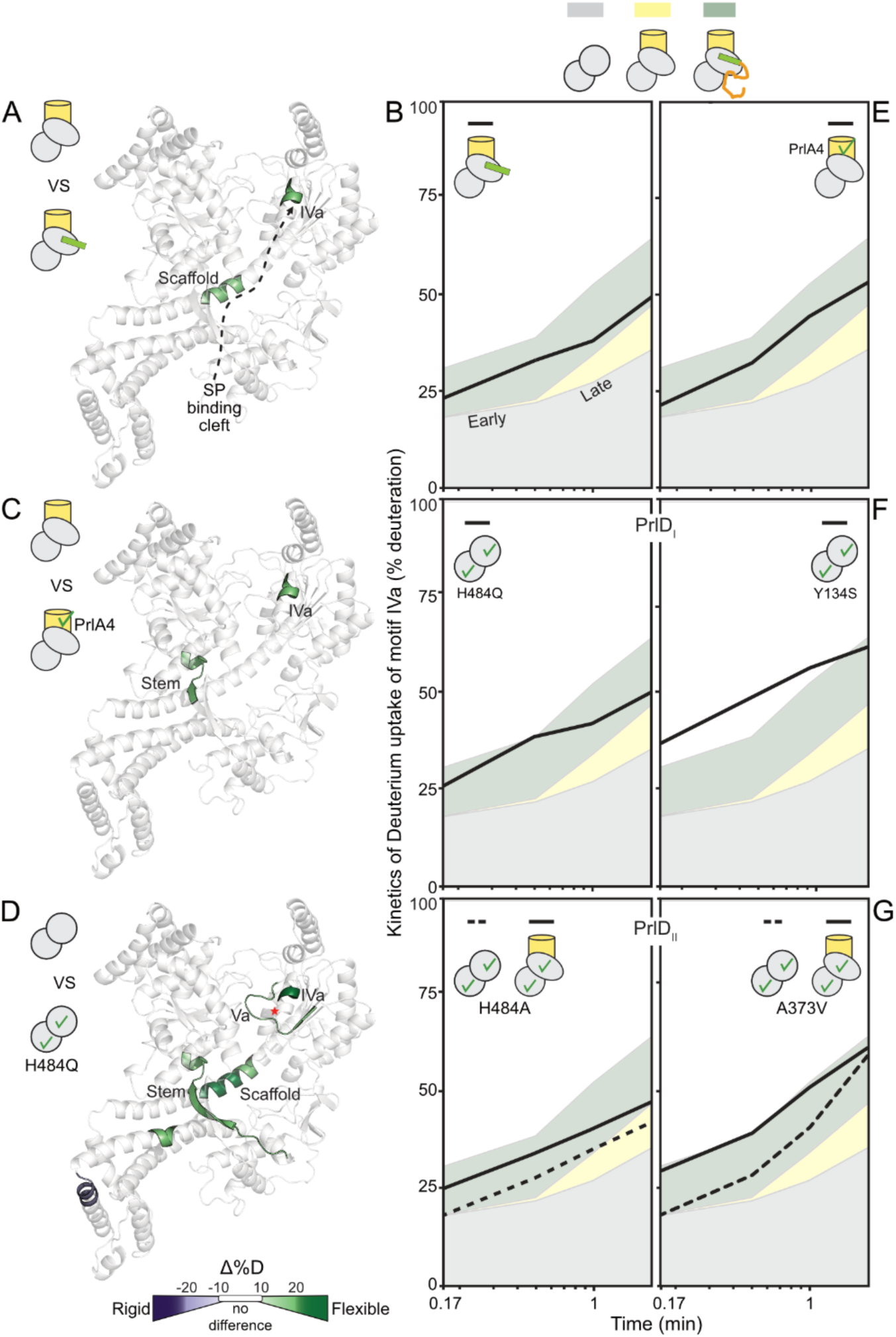
Signal peptides trigger the translocase by enhancing gate2 dynamics. **A.** Long range signal peptide effect shown schematically (dashed arrow) on the local dynamics of channel-primed SecA_2_. Regions showing differential D-uptake in SecYEG:SecA_2_:signal peptide compared to SecYEG:SecA_2_ are mapped onto the structure of a single SecA protomer, as indicated. Only increased dynamics were observed (green). **B., E., F. and G.** D-uptake kinetic plots of a motif IVa peptide (aa488-501, as in Fig.1E) under the indicated conditions are compared with the kinetics of the same peptide from SecA_2_ (grey), SecYEG:SecA_2_ (yellow) and SecYEG:SecA_2_:preprotein (green) in the absence of ADP (see also Fig. S2A, left). Minor differences were seen between the ADP bound (Fig.1E) and free (Fig. S2A) states. Selected data points (10 s, 30 s, 1 m, 2 m labeling times) focus on the kinetic regime with maximum differences; SD values (<2%) have been removed. *n = 3*. **B.** D-uptake kinetics of motif IVa peptide in SecYEG:SecA_2_:signal peptide state (green line). **C.**-**D.** Local dynamics of SecY_PrlA4_EG:SecA_2_ (test) compared to those of SecYEG:SecA_2_ (control)(**C**); of SecA(H484Q)_2_ (test) compared to those of SecA_2_ (control) (**D**); as in B. Red asterisk: H484Q. **E.**-**G.** D-uptake kinetics of motif IVa peptide (as in B) in SecA_2_:SecY_PrlA4_EG translocase (**E**) or the indicated PrlD_typeI_ (**F**) and PrlD_typeII_ (**G**) mutants in solution.

Therefore, the signal peptide caused measurable but minor (compared to those of the preprotein) rearrangements on the conformational landscape of SecA, primarily at motif IVa of the peripheral gate2. This may be essential for ‘triggering’ (Fig. 1A.III)(Gouridis et al., 2009).

### Signal peptide-induced translocase triggering occurs via motif IVa dynamics

To better understand how signal peptides control translocase dynamics via motif IVa we used Prl (*pr*otein *l*ocalization) mutants in SecA (PrlD) or SecY (PrlA). These gain-of-function mutants are informative because they can secrete clients devoid of signal peptides (Fig. S2C) (Flower et al., 1994; Huie and Silhavy, 1995), and therefore are structural mimics of the signal peptide-induced states. They achieve this because they exist constitutively in the triggered conformation [Fig. 1A.III; Fig. S2D, lanes 2-5; (Gouridis et al., 2009)]. In fact, some of them are triggered even in the absence of the channel (Fig. S2D, lanes 4 and 5; hereafter SecA_PrlDI_).

Remarkably, three SecA_PrlD_ hotspot residues line both sides of gate2: H484 and A488 in motif IVa, juxtapose Y134 of motif Ia [Fig. S2C; (Huie and Silhavy, 1995)]. Other hotspot residues lie in adjacent motifs, e.g. A507 in motif Va.

Minor side chain alterations in either Y134 or H484 mimic the binary effect of channel plus signal peptide binding, in the absence of either. We dissected the two ligand effects on H484, by screening mutant derivatives. SecA(H484A) also displays a Prl phenotype (Fig. S2E, lane 6). However, unlike SecA(H484Q), its triggering required prior channel binding (Fig. S2D, lanes 6-7; hereafter SecA_PrlDII_).

We characterized the local dynamics of these mutants. Wild type SecA_2_ bound to SecYEG or to SecY_PrlA4_EG exhibited similar dynamics (Krishnamurthy et al., 2021) but the latter displayed additionally elevated dynamics in motif IVa and the Stem (Fig. 2C). Similarly, compared to SecA_2_, SecA(H484Q)_2_ exhibited elevated motif IVa, Stem and Scaffold dynamics in the absence of channel or preprotein (Fig. 2D).

Remarkably, in time-dependent dynamics analysis of Motif IVa, SecA_2_:SecY_PrlA4_EG (Fig. 2E) and the SecA_PrlDI_ mutants alone (Fig. 2F] all displayed elevated early phase dynamics, similar to those driven by preprotein or signal peptide in the wildtype (Fig. 2B). SecA_PrlDII_ mutants also displayed moderately elevated motif IVa dynamics (Fig. 2G; dashed line) that were increased significantly after channel addition (solid line) thus, explaining their channel-dependence for triggering.

Fine modulation of the complex dynamics at motif IVa of gate2 is an essential aspect of signal peptide-mediated translocase triggering.

### Signal peptides promote closing of the preprotein clamp

Signal peptide binding at PBD (Gelis et al., 2007), influences motif IVa dynamics, over 4 nm away, and triggers the translocase. To explain how, we hypothesized that signal peptides might influence PBD rotation around its Stem (Krishnamurthy et al., 2021; Vandenberk et al., 2019). We probed such domain motions using smFRET. Fluorophores on PBD and NBD2, that form the preprotein clamp (Fig. S3A), can monitor these domains coming close or apart [Fig. S3B; yielding high/low FRET respectively; (Krishnamurthy et al., 2021; Vandenberk et al., 2019)].

In SecA_2_:SecYEG the PBD of the active protomer samples all three states, with a preference for the Wide-open [Fig. 3A, lanes 1-3; S3B.IIa; (Krishnamurthy et al., 2021)]. Preprotein binding led to clamp closing (i.e., Open plus Closed states) in 98% of the active protomers (lanes 4-6), irrespective of ADP presence (Fig. S3B.III). Signal peptides alone can replicate this in 85% of the active protomers (Fig. 3A, lanes 7-9). SecA_2_:SecY_PrlA4_EG that is already triggered promotes clamp closing in the absence of preprotein or signal peptide (lanes 10-12). Signal peptide-driven clamp closing (Fig. 3A, lanes 7-9) is not accompanied by any detectable secondary structure/flexibility changes inside the PBD or the nucleotide cleft (Fig. 2A). This mainly rigid body motion seems rather uncoupled from stimulating nucleotide turnovers in the helicase motor (see below). In freely diffusing wild type SecA_2_ the clamp equilibrium is maintained at the Wide-open state [Fig. 3B, lanes 1-3; (Krishnamurthy et al., 2021)] and signal peptides do not close it (Fig. S3B.I).

**Figure 3:**
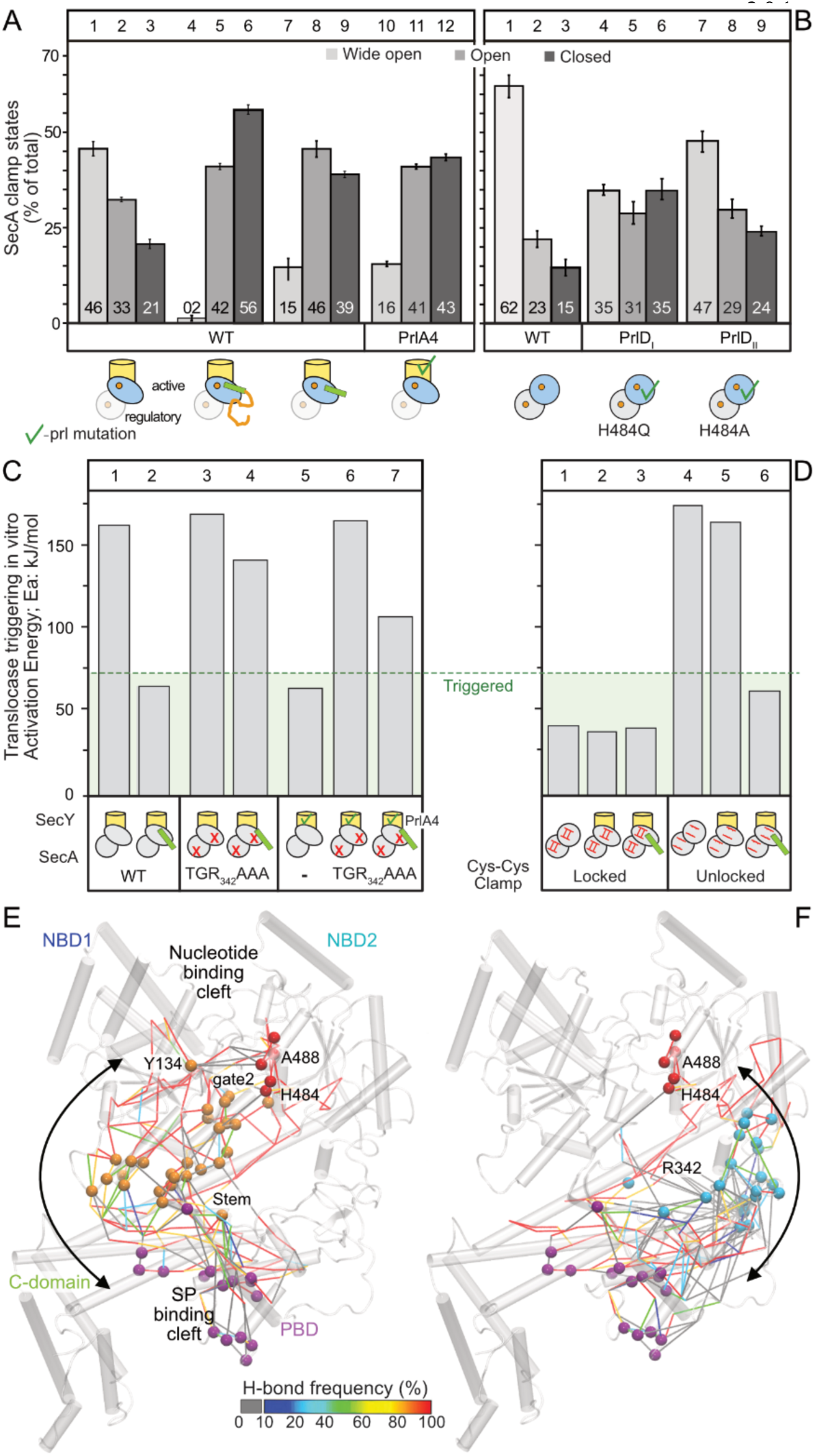
Clamp closing is a major conformational event towards translocase triggering. **A-B**.Distribution of the preprotein clamp states of Channel-bound SecA_2_ states (**A**) and free SecA_2_ states (**B**), determined by freely diffusing confocal smFRET as described (Krishnamurthy et al., 2021). His-SecA (50-100 pM), stochastically labelled at V280C_PBD_/L464C_NBD2_ with Alexa 555 and Alexa 647 (blue circle) was allowed to dimerize with excess cold SecA (1 μM; grey circle) resulting to SecA_2_ with a single fluorescent protomer. Gaussian distributions, fitted to FRET histograms and quantified, correlate to the wide-open, open and closed clamp states (Fig. S3A-B). Under channel-bound conditions, only data for the active, channel bound protomer (blue oval) are shown (Krishnamurthy et al., 2021). *n ≥ 3*; mean (} SEM). **C.** and **D.** Activation energy (*E_a_*) for wild type SecA, SecA(TGR_342_AAA) (**C**) and a double cysteine SecA derivative (**D**) under the indicated conditions. Oxidized/locked Closed, or reduced/unlocked Open clamp as indicated. **E.** and **F.** H-bond pathways connecting motif IVa (red) to the signal peptide cleft (purple) through either the Stem (**E**; orange) or the PBD-NBD2 interface (**F**; cyan), derived from graph analysis of MD simulations of ecSecA_2VDA_ with an open gate2 (Krishnamurthy et al., 2021).

In contrast, in diffusing, spontaneously triggered SecA_PrlDI,_ clamp equilibria shift towards closed states in the absence of channel and signal peptides (Fig. 3B, lanes 4-6) and less so in SecA_PrlDII_ mutants that require the channel for triggering (lanes 7-9; Fig. S3B.IV.b-c; Fig. S3C).

Our results raised the possibility that the direct physical interaction of NBD2 with PBD that is promoted in the Closed state might be functionally important. To test this, we mutated the highly conserved 3β-tip of PBD that binds to NBD2 to close the clamp [Fig. S3A; (Krishnamurthy et al., 2021)]. The generated SecA(TGR_342_AAA) failed to become triggered by signal peptide (Fig. 3C, lane 4) or SecY_PrlA4_ (lane 6) or their combination (lane 7). SecA(TGR_342_AAA) binds to channel/preproteins (Fig. S4A) yet, fails to stimulate its ATPase or secrete *in vitro* or complement *in vivo* function (Fig. S4B-D).

To further probe the importance of the closed clamp state we locked it closed through engineered disulfides (Chatzi et al., 2017; Sardis et al., 2017) and tested the functional consequences. The SecA_locked closed_ was permanently triggered, independently of channel or preprotein (Fig. 3D, lanes 1-3), akin to SecA_PrlDI_ mutants (Fig. S2C, lanes 4-5). Reduction of the disulfide reinstated a ‘channel plus preprotein’ requirement for triggering (Fig. 3D, lanes 4-6).

Signal peptide-driven clamp closing and increased motif IVa dynamics underlie translocase triggering.

### The signal-peptide cleft cross-talks to motif IVa via two main H-bond pathways

To determine how signal-peptide driven clamp closing might allow the signal peptide binding cleft to cross-talk with motif IVa, we determined the H-bonding networks, including water-mediated bridging, between the two allosterically connected sites. In all simulations of *ec*SecA (monomeric or dimeric), motif IVa (Fig. 3E-F, red spheres; Table S2) was interconnected within a local H-bond network that could extend to involve most of SecA’s residues (Krishnamurthy et al., 2021). Graph analysis determined the most frequently visited/shortest possible H-bond pathways that could be potentially altered along the reaction coordinate of SecA through which, the signal peptide cleft in PBD (purple spheres) could communicate with motif IVa. Two main routes were proposed by this analysis. One was via the NBD2_Joint_/PBD_Bulb_ interface of the closed clamp (Fig. 3F, cyan spheres) that was experimentally tested above. The other was via the PBD_Stem_/a8 interface which binds mature domains and interconnects to the second half of gate2 (Y134; Fig. 3E, orange spheres). This second route was tested below.

### The signal-peptide cleft cross-talks to motif IVa through the Stem/α8 interface

The Stem/*α*8 interface that binds mature domains (Chatzi et al., 2017), undergoes significant conformational changes during the transition from the Open to the Closed state [Fig. 4A; S3A; (Krishnamurthy et al., 2021)]. These might alter the local hydrophobic interactions between the Stem *β* strands and *α*8, as this interface is pried open. The Stem/α8 interface extends to a three strand β-sheet with the highly dynamic β6_Stem_ and β24_C-tail_ and involves L187*α*8 that packs against A373 of *β*12_Stem_ (Fig. 4A).

**Fig. 4.**
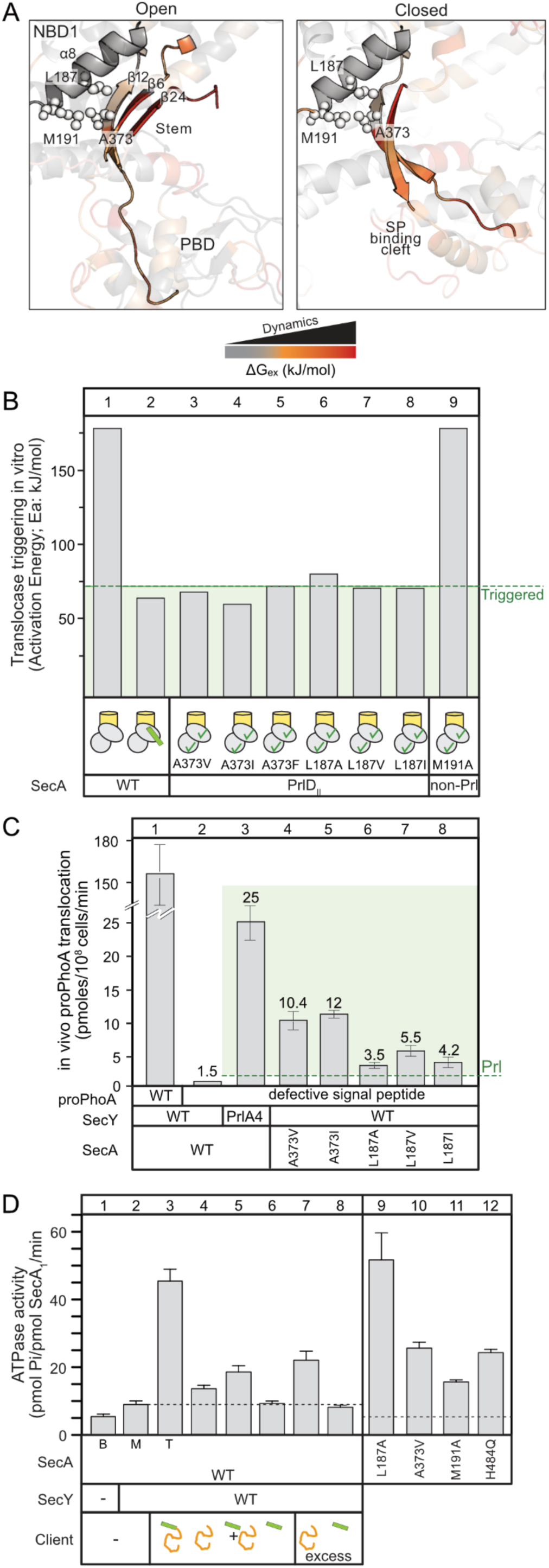
Mature domain-driven ADP release and ATP turnovers. **A.** Structure and residues at the Stem/*α*8 region including β24_C-tail_. ΔG_ex_ values are shown for the SecYEG:SecA_2_:preprotein state in the Open (middle; *ec*SecA_2VDA_) and Closed (right; *ec*SecA_2VDA_-MD model;(Krishnamurthy et al., 2021)) clamp conformation. Structures are aligned based on NBD1. **B.** Activation energy (*E_a_*) for the indicated Stem mutants with SecA_PrlDII_ phenotype in channel-primed states, compared to wild type translocase (as in Fig. 3B). ***C.*** *In vivo* translocation of proPhoA or of a defective signal peptide derivative [pro(L8Q)PhoA] by the indicated translocases. Secreted phosphatase units were converted to protein mass, as described (Gouridis et al., 2010). *n= 6*; mean values (± SEM). **D.** The ATPase activity of freely diffusing (basal; B; 0.4µM SecA or the indicated derivatives), SecYEG-bound (membrane; M; 1 µM SecY) and translocating SecA (T; SecY plus 9 µM proPhoACys_-_) was determined as described (Gouridis et al., 2010). Signal peptide (30 µM; excess: 60 µM), mature domain (PhoAcys-; 20 µM; excess: 40 µM). *n= 3-6*; mean values (± SEM).

To strengthen or weaken hydrophobic packing at the Stem/*α*8 interface we substituted A373_Stem_ with large hydrophobic residues (V, I and F) and L187*α*_8_ with V, I or A. All derivatives displayed Prl phenotypes *in vitro* (of Type II)(Fig. 4B, lanes 3-8) and *in vivo* (Fig. 4C, lanes 6-8) with L187A the weakest one. A373V was previously known as the only Prl outside the nucleotide binding cleft (Flower et al., 1994; Huie and Silhavy, 1995). In contrast, mutating M191 to A, located one turn after L187 at the end of *α*8 and of the Stem/*α*8 interface (Fig. 4A), did not yield a Prl phenotype (Fig. 4B, lane 9). All mutant derivatives were functional *in vivo* (Fig. S4E).

We concluded that through this route signal peptides alter hydrophobic packing at the mature domain binding patch on the Stem/α8 interface.

### Binding of mature domains drives ADP release and ATP turnover

Preprotein mature domains bind to the Stem/*α*8 interface [(Chatzi et al., 2017); Fig. S1B, orange surface] and when covalently associated to signal peptide, stimulate ADP release (Fig. 1B) and multiple ATP turnovers on the translocase (Fig. 4D, lane 3; (Karamanou et al., 2007). Even mature domains alone (lane 4) or together with signal peptides added *in trans* (lane 5) or alone in excess (lane 7) drive some measurable ATPase stimulation (1.5, 2 and 3-fold, respectively), while signal peptide alone (lane 6), even in excess (lane 8), does not.

Mature domains cause minor increased flexibility in the local dynamics of the helicase motor of SecYEG:SecA_2_:ADP that is not imrpoved with the simultaneous addition of signal peptides *in trans* (Fig. S4F-G). This explains their inability for full ATPase stimulation when the two moieties are not covalently connected.

Based on the above, we hypothesized that signal-peptide induced effects could positively contribute through conformationally optimizing the Stem/*α*8 interface for mature domains to bind and stimulate ATP turnovers. To directly test the possible role of this region in regulating ATP turnover, we screened the Stem/*α*8 interface for mutant derivatives that exhibit high basal ATPase activity mimicking the mature domain-bound state. Prl mutants (L187A and A373V located at Stem/*α*8; Fig. 4B) displayed elevated ATPase activity compared to free SecA_2_ (Fig. 4D, lanes 9-10). In contrast, mutant derivatives of residues in the back face of *α*8 did not and had compromised function (Fig. S4H-J).

Clearly, the signal peptide conformational effect on Stem/*α*8 and the ATPase activity of the motor are tightly coupled. We disentangled the two effects by characterizing the mature domain binding site residue (M191; Fig. 4A). Freely diffusing SecA(M191A) displays ∼3-fold elevated basal ATPase (Fig. 4D, lane 11), that was hyper-stimulated ATPase under translocation conditions (Fig. S4K), but neither displayed a Prl phenotype (Fig. 4B, lane 9), nor showed measurable changes in the dynamics of motif IVa (Table S1). Residues like M191 at the Stem/*α*8 interface appear critical in allowing mature domains to control the ADP release cycle of the helicase motor.

Our results show that signal peptides and mature domains have functionally inter-connected but divergent roles in activating the translocase, that converge at the Stem/α8 interface. This interface is a conformational hub that couples the rate limiting ADP release cycle (Fak et al., 2004; Sianidis et al., 2001) and the motif IVa dynamics in the helicase motor to conformational cues sent by the signal peptide upon closing the clamp and promoting mature domain binding. Only legitimate secretory clients that satisfy all these relationships or Prl mimics, induce the ATPase activity of the translocase (Robson et al., 2009).

### Nucleotides finely control the intrinsic dynamics of SecA

Co-ordinated docking of signal peptides and mature domains causes loss of ADP from the SecA helicase motor, allowing initiation of a fresh ATPase cycle. We hypothesized that a core role of the nucleotide cycle may be to modulate the intrinsic dynamics of SecYEG:SecA_2_. To examine this, we used nucleotide analogs that represent different stages of the ATPase cycle (Fig. S5A). Nucleotides bind in a positively charged cleft between NBD1 and 2 (Fig. S5B) and make contacts with residues from both domains, mostly with NBD1; β and *γ*-phosphates also with NBD2 (Krishnamurthy et al., 2021; Papanikolau et al., 2007). Signal peptides and mature domains on the other hand, bind on negatively charged surfaces that line the clamp and signal peptide binding cleft (Fig. S5B). Using HDX-MS we monitored the dynamics of the quiescent ADP state (Krishnamurthy et al., 2021), pre-formed at 2mM (10_4_-fold excess over K_d_) to overcome the ADP release reaction, the apoprotein (i.e. cleft emptied due to preprotein binding; Fig. 1B), the ATP-bound but not yet hydrolyzed state (mimicked by the non-hydrolyzable analogue AMP-PNP) and the ATP hydrolysis transition state intermediate [mimicked by ADP:BeF_X_ (Zimmer et al., 2008)]. We compared the dynamics of the SecYEG:SecA_2_:ADP (Fig. 5 and Fig. S5C; reference state, top pictogram) to all other nucleotide states (test states, bottom pictograms).

**Figure 5.**
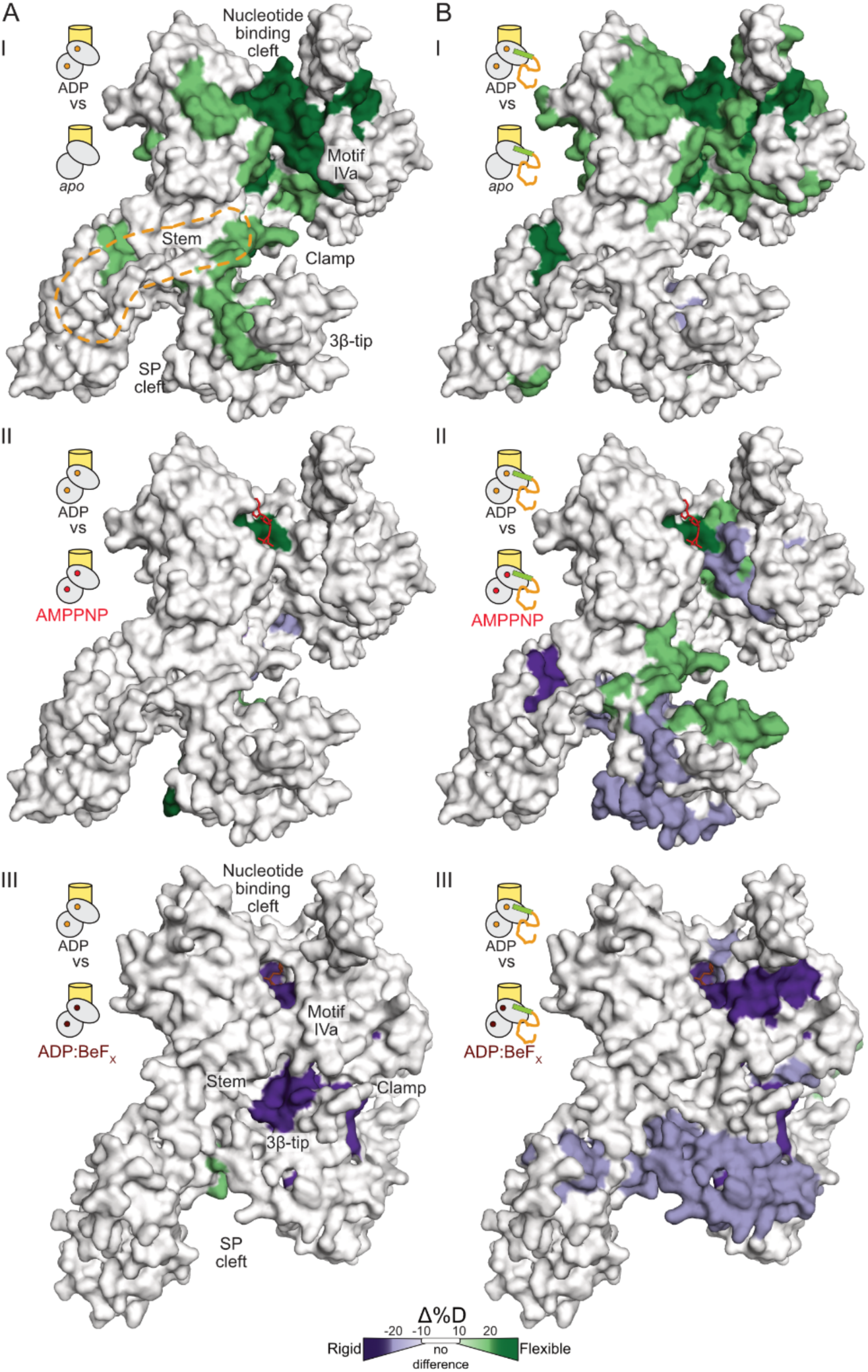
Nucleotide states and preproteins drive distinct translocase conformational motions. **A.** and **B.** The local dynamics of the SecYEG:SecA_2_ translocase at the indicated nucleotide state was compared to those of the SecYEG:SecA_2_:ADP one (‘reference’), in the absence (**A**) or presence of preprotein (**B**). Regions showing differential Duptake in (I) apo, (II) AMPPNP-bound and (III) ADP:BeFX-bound states (“test”) were colored mapped on a SecA protomer visualized by surface representation. PatchA is indicated in A.I by orange dashed line. Apo (I) and the AMPPNP bound (II) states are mapped on the open clamp PDB: 2VDA structure; the DP:BeFX bound state (III) is shown on the closed-flipped clamp PDB: 3DIN structure. ADP: orange circle, AMPPNP: red circle, ADP:BeFX: brown circle

SecA_2_:ADP in solution is auto-inhibited, until primed by channel binding (Krishnamurthy et al., 2021). This primed translocase is conformationally stabilized, binds preproteins with high affinity but does not exhibit elevated ATPase activity until preproteins bind and cause ADP release (Fig. 1B). Emptying the cleft of ADP causes widespread elevated dynamics in the helicase motor and more localized ones in the Stem, Scaffold, IRA1 and PBD [Fig. 5A.I; (Krishnamurthy et al., 2021)]. SecYEG:SecA_2_:AMPPNP showed minor differences in dynamics compared to SecYEG:SecA_2_:ADP (Fig. 5A.II; S5C.II). ADP:BeF_X_ caused additional stabilization at the motor (motifs I, IVa, V/Va) and the PBD clamp region (Stem, *α*13 and the 3β-tip_PBD_ that binds NBD2) (Fig. 5A.III). Motif Va contains the R509 arginine finger, crucial in γ-phosphate recognition and regulation of motor conformational states (Keramisanou et al., 2006). These results are also consistent with the crystal structure of SecYEG:SecA:ADP:BeF_x_ [Fig. S6A; (Zimmer et al., 2008)]. In this “closed-flipped” state the PBD_bulb_ and NBD2 home into each other and the PBD_3β-tip_ also flips towards NBD2_motifIVa_, salt-bridging the essential R342 with E487 (Fig. S6B-C Movie S1). Conversion to the ADP state, reverses rigidification through minor local changes (Fig. S6D.VIII). Importantly, some of these intra-protomeric changes, such as the enhanced ADP-driven dynamics, extend to enhancing the dynamics of the SecA dimerization interface (Fig. S5C.I), suggesting preparatory steps towards dimer dissociation (Gouridis et al., 2013).

The effects described above are only specific to SecA that has been primed by binding to SecY. Soluble SecA_2_:ADP:BeF_X_ only shows weak contacts (Table S1) in the helicase motor with overall higher dynamics than SecA_2_:ADP (Fig. S6D.IV; conversion from ADP:BeF_X_ to ADP results in decreased dynamics in the ADP state).

The Q motif that tightly binds the immutable adenine ring (Fig. S1E), showed negligible dynamics in the presence of nucleotide (Krishnamurthy et al., 2021). This presumably explains the similar high affinities of ATP derivatives for the helicase motor. It is the mutable *γ*-phosphate ends of nucleotides like AMPPNP and ADP:BeF_x_ that bind to NBD2 and stabilize different motor conformations, while ADP does not (Fig. S1E; S5A). Thus, missing the *γ*-phosphate contacts, weakens the NBD2-ADP association, allowing higher mobility of the nucleotide inside the pocket and increased Motif I dynamics (Fig. S6D.VIII). These data can explain how, despite their minor chemical differences, nucleotides display major differences in stabilizing unique conformational states of SecA by exploiting its intrinsic dynamics.

Therefore, nucleotides exploit the intrinsic dynamics of SecA by promoting multiple but minor local structural dynamics changes, largely driven by interactions of their *γ*-phosphate end with NBD2. None of the nucleotide effects described above alter the domain motions of the PBD clamp significantly (Fig. S6E).

### Preproteins regulate nucleotide-controlled dynamics in channel-bound SecA

Next, we probed how preproteins might exploit the intrinsic, nucleotide regulated conformational dynamics of the primed translocase leading to active translocation. For this, we followed translocase dynamics of SecYEG:SecA_2_:different nucleotide (as in Fig. 5A) but in the presence of the proPhoA_1-122_ secretory client (15 µM; >50 fold over K_d_; Fig. 5B, Fig. S5D).

Preprotein-driven ADP release from the translocase (Fig. 1B), led to increased dynamics in the helicase motor of the resulting translocase apoprotein (Fig 5B.I, Fig. S6C), similar to those seen without the preprotein (Fig. 5A.I). AMPPNP binding to the preprotein-bound translocase, caused significant differences compared to those of ADP binding (Fig. 5B.II): enhanced dynamics in the helicase motifs of the nucleotide binding cleft (I, Ic, III and VI; Fig. S1E) and inside of the clamp (3β-tip and β24 of the C-tail; Fig. 4A) and decreased dynamics at the signal peptide binding cleft (Stem, α10, α13). Contrasting dynamics on either side of the PBD suggest that ATP binding has divergent effects on the signal peptide and mature domain segments of the client. ADP:BeF_X_ reduced dynamics in all of the helicase motifs of NBD2 without affecting NBD1 and rigidified wide areas of the clamp (Stem, α13 and 3β-tip, Joint, Scaffold; Fig. 5B.III). While these local dynamics changes are coincident with closing of the preprotein clamp, the latter occurs independently of nucleotide and is driven by signal peptide binding (Fig. S3B.III). The important mechanistic implication of these observations is that preprotein clients physically couple the otherwise unconnected nucleotide-regulated helicase motor dynamics to clamp motions.

Our results provided insights into conformational processes that are nucleotide or/and preprotein-driven (Fig. 5). Nucleotides intimately regulate the dynamics of the helicase motor, with minor effects on any preprotein binding region. In the presence of preproteins not only the nucleotide effects at the motor persist, but the dynamics of the preprotein binding sites now become modulated by the nucleotide state. Clearly, preprotein binding on multiple SecA locations (Chatzi et al., 2017; Sardis et al., 2017) requires concerted motions of various translocase regions to achieve translocation.

### Locally frustrated regions in the SecA clamp allow client promiscuity

One of the intriguing aspects of the translocase is that it handles hundreds of non-folded clients with dissimilar sequences that are all expected to bind to the same preprotein binding sites on SecA. Such promiscuity is a fundamental universal feature of chaperones. In the absence of conserved linear features, non-folded clients are thought to be recognized through their frustrated regions (He and Hiller, 2019; He et al., 2016) by frustrated local structural elements in chaperones [(Hiller, 2019; Morgado et al., 2017) Srinivasu et al., submitted]. Frustrated structural elements sample multiple degenerate conformations, that allow chaperones to mold its binding interfaces in order to recognize and bind a wide range of clients. As frustrated regions commonly display elevated dynamics [(Hiller, 2019; Morgado et al., 2017)(Srinivasu et al., submitted)], reminiscent of those seen around the translocase clamp, we hypothesized that SecA may use similar mechanisms. To test this, we compared the dynamic islands determined experimentally by HDX-MS [Fig. 6A; regions with low ΔG_ex_ values; orange/red regions)(Krishnamurthy et al., 2021)] to frustrated regions predicted from the *ec*SecA_2VDA_ structure using the tool frustratometer (Parra et al., 2016)(Fig. 6B). Interestingly, most frustrated inter-residue contacts (Fig. 6B; green lines) and the experimentally determined dynamic islands (e.g. the Joint and motif IVa in NBD2; prong 1, 3β-tip (prong 2) of PBD and the tip of IRA1) closely overlapped. Clamp closing due to signal peptide binding (Fig. 3A, lanes 7-12) would allow the 3β-tip, prong 2 and Joint interact to form a contiguous region of frustration (Fig. S7A.II) that could trap preproteins by forming local favourable interactions with the client. In the closed-flipped state of the ATP hydrolysis transition state (ADP:BeF_X_)[Fig. S7B; (Zimmer et al., 2008)] two parallel regions of frustration become evident that together with the closed clamp could enclose/can interact with the preprotein chain as it enters the channel.

**Figure 6.**
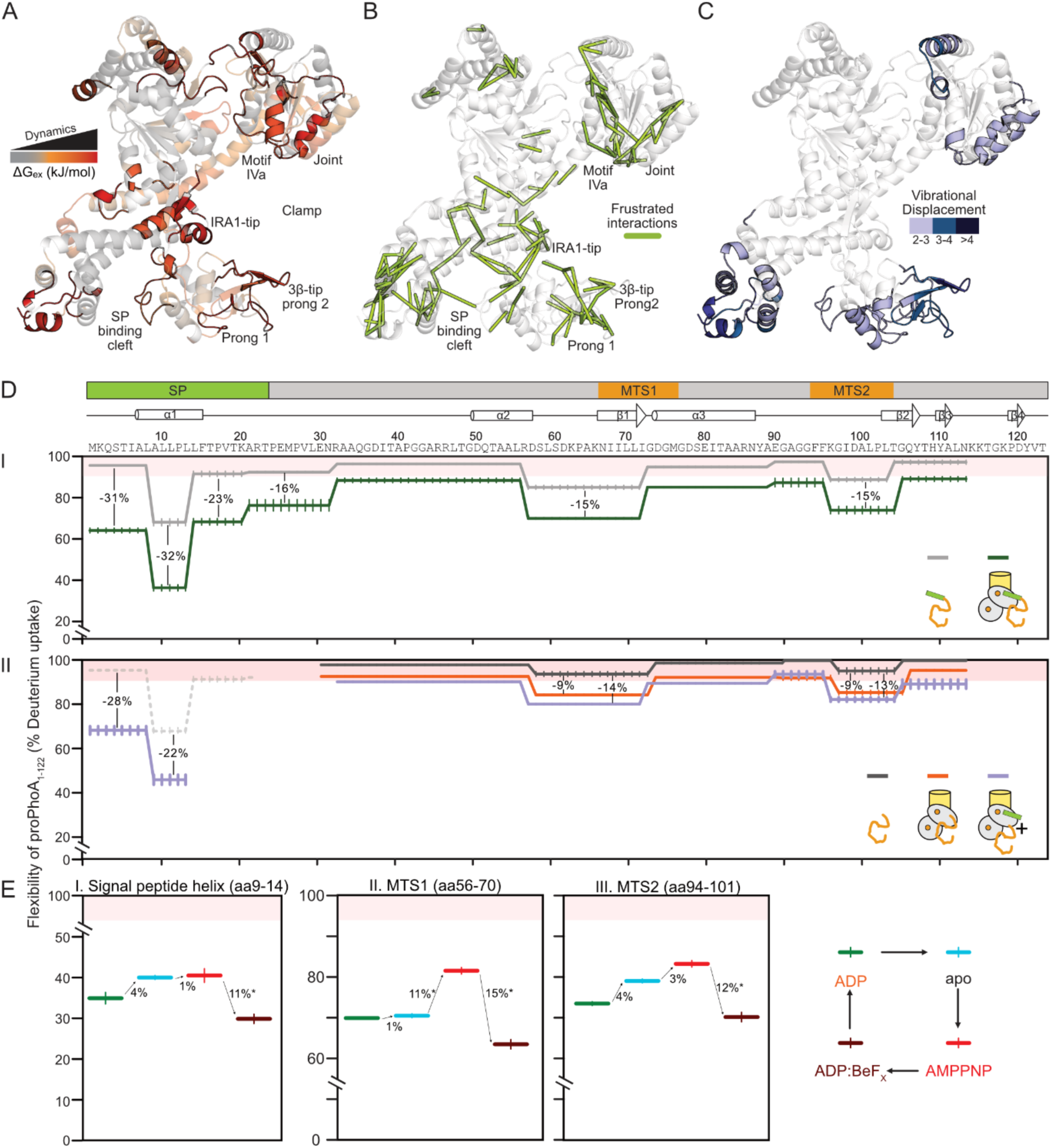
Translocase binds and regulates islands of preprotein dynamics. **A.** ΔG_ex_ values were calculated for free SecA_2_ and mapped onto the open clamp structure (PDB ID:2VDA). Highly flexible regions (i.e. ΔG_ex_= 11-16 kJ/mol) are indicated with red hues. **B.** SecA regions of frustration (green) derived from the Frustratometer server (Parra et al., 2016) using PDB ID:2VDA as input. **C.** Total displacement of normal modes 7-12 (unweighted sum) derived from PDB ID: 2VDA are mapped in blue shades onto the SecA structure. See detailed analysis in Fig. S7C. **D.** HDX-MS-derived flexibility map of proPhoA_1-122_ (I) and PhoA_23-122_ (II) shows the absolute dynamics (in % D-uptake) of the preprotein at residue level (from N to C terminal), in the indicated states. Higher % D-uptake values correlate with increased flexibility, residues showing >90% flexibility values are considered hyper-flexible. I. The flexibility of free (grey) and SecYEG:SecA_s_:ADP bound proPhoA_1-122_ (green) are compared. II. The flexibility of free (black) and SecYEG:SecA_2_:ADP bound PhoA_23-122_ (orange) are compared. The effect of the addition of signal peptide in trans on the dynamics of SecYEG:SecA_2_:ADP:PhoA_23-122_ is shown in purple. As a control, the flexibility of the signal peptide as part of free proPhoA_1-122_ (as in I; dashed grey) is shown. Differences greater than 10% are considered significant. Residues are aligned with known secondary structural features of native PhoA. The position of signal peptide and MTS (mature domain targeting signals) 1 and 2 are shown. *n = 3*, SD values are represented as vertical lines. SD values <1 % fall within the width of the line and hence are omitted. **E.** Flexibility map of three selected regions of proPhoA_1-122_ bound to channel-primed translocase in the presence of various nucleotides. The nucleotide cycle (right pictogram) follows the translocase from ADP bound state (green; ground state), followed by ADP release to apo state (light blue), ATP binding state (mimicked by AMPPNP; red) and ATP hydrolysis transition state (mimicked by ADP:BeFX; brown). The flexibility of (I) signal peptide, (II) MTS1 and (III) MTS2 of the proPhoA_1-122_ is monitored as the translocase is going through the nucleotide cycle, with differences in % D-uptake between nucleotide states quantified.ss

We additionally probed the inherent dynamics of SecA using an orthogonal biophysical tool, Normal Mode Analysis (NMA) (on PDB: 2VDA). NMA provides a mathematical description of atomic vibrational motions and protein flexibility (Bahar et al., 2010; Kovacs et al., 2004). In this method, Cα atoms (modeled as point masses) are considered to be connected by springs that represent interatomic force fields. The resulting model generates a set of “normal modes”, where all Cα atoms are oscillating with the same frequency. The lowest frequency normal modes contribute the most to domain dynamics within a protein and the associated Cα displacement can be calculated [Fig. S7C; (Hinsen, 1998; Tiwari et al., 2014)]. Motif IVa and the Joint in NBD2, 3β-tip in PBD and the signal peptide binding cleft are regions that show maximum displacement during vibrational motions (Fig. 6C; shades of blue) and practically coincide with the experimentally determined dynamic islands and the frustrated regions.

Taken together, our data suggest that altered dynamics in these regions is a direct ramification of nucleotide/client mediated modulation and that non-folded clients would recognize them promiscuously forming local less frustrated interactions to achieve the coupling of the dynamics of these regions to the ATPase cycle.

### Signal peptide-driven closing of the clamp allosterically enhances mature domain binding

The data above revealed a nucleotide-regulated structural dynamics framework in SecA that provides for multi-valent localized, transient interactions with the non-folded clients. To decipher how these dynamics are translated into translocation steps for the translocating client, we developed an HDX-MS-based assay to directly monitor the dynamics of the proPhoA_1-122_ client (Fig. S7D). ProPhoA_1-122_, contains the three necessary and sufficient elements for high-affinity binding to the translocase and secretion: a signal peptide and the first two mature domain targeting signals of ProPhoA (MTS1 and 2; Fig. 6D, top; (Chatzi et al., 2017)] and allowed translocation to be dissected away from folding processes (Tsirigotaki et al., 2018).

proPhoA_1-122_ was diluted from chaotrope into deuterated buffer, pepsinized, its peptides analyzed by mass spectrometry and their % D uptake was determined. 90-100 % D-uptake values signify extensive dynamics/lack of stable secondary structure [Fig. 6D.I, pink area; (Tsirigotaki et al., 2017b)]; segments of folded proteins with ordered secondary structure typically exhibit 20-40% D-uptake values.

proPhoA_1-122_ alone, in solution, at 37_o_C, displays high % D uptake values (Fig. 6D.I, grey), consistent with a protein remaining extensively non-folded under these conditions. Three islands of proPhoA_1-122_ sequence show slightly lower % D uptake values suggestive of some stabilization of the backbone amides. This is likely due to hydrogen-bonding upon transient acquisition of local secondary structure and indeed these regions overlap, wholy or partly, with secondary structural elements in the natively folded PhoA (Fig. 6D, top, 2ndary structure map). These sequences include: a. the helical hydrophobic core of the signal peptide (aa 9-13) and its downstream region (∼17 residues), including the early mature region “rheostat” that controls PhoA folding (Sardis et al., 2017); b. the hydrophobic core of MTS1 (aa 68-72) and its upstream more polar stretch (aa 56-68). c. the hydrophobic core of MTS2 (aa 94-102).

Dilution of proPhoA_1-122_ from chaotrope into solution containing SecYEG:SecA_2_, at 37°C, reduced % D uptake significantly in all of the wider signal peptide region (aa 1-33; including the positively charged N-region and the downstream rheostat but more prominently in the signal peptide helix) but also substantially in the MTS1 and 2 islands (Fig. 6D.I, green line). This is indicative of direct SecA binding on these segments and was also seen on peptide arrays (Chatzi et al., 2017). Rigidification likely reflects more stable H-bonding either within the secondary structure elements (e..g. signal peptide *α*-helix; *α*2 in MTS1) or externally with SecA. In contrast, the linkers connecting these three regions remain highly unstructured (>85 % D uptake). Monitoring these islands of differential localized highly dynamic translocase binding observed on the client chain is a unique power of this HDX-MS assay.

Covalent connection between signal peptide and mature domain was important for optimal binding to SecA. Mature domain alone bound to SecYEG:SecA_2_ more weakly than did the preprotein (Fig. 6D.II, compare black to orange and to Fig. 6D.I). This binding was further stabilized by addition of signal peptide *in trans* (Fig. 6D.II, purple line) or use of SecY_PrlA4_:SecA_2_ (Fig. S7E, purple), but never reaching the extent of binding seen with preprotein.

These data provided a direct explanation for the importance of the covalent connection in the preprotein moieties seen above in achieving maximal translocase interaction (Fig. 1B; 1E; S2B; 4D). They also suggested that SecA and the three client islands inter-communicate allosterically.

### The nucleotide cycle selectively alters SecA interaction with the signal peptide and mature domain islands

To dissect preprotein dynamics during the ATPase cycle of the translocase we monitored the preprotein interaction with each of the four unique translocase:nucleotide/analogue conformations (Fig. 6E, right), specifically focusing on the dynamics of the three main islands that bind SecA: the hydrophobic helix of the signal peptide, MTS1 and 2 (Fig. 6EI-III).

The signal peptide shows slightly tighter binding when transitioning from the ADP-bound to the apoprotein to the AMPPNP-bound translocase but becomes significantly rigidified on the ADP:BeF_X_-bound translocase (Fig. 6E.I, red to brown). MTS1 dynamics are unchanged when transitioning from the ADP-bound to the apo translocase but become significantly increased in the AMPPNP state (Fig. 6E.II, blue to red) and significantly decreased in the ADP:BeF_X_ state (red to brown). For MTS2, as the ATPase cycle progresses from ADP-bound to apo to AMPPNP state, its dynamics increase incrementally (Fig. 6E.III, green to blue to red) but, they decrease significantly in the ADP:BeF_X_ state (red to brown).

Taken together, these results showed that in the ADP and apo stages of the translocation ATPase cycle, the SecA subunit binds potentially all three elements of the client. Upon ATP binding SecA loosens its grip on the mature part of the chain, as evidenced by the overall increased dynamics in the AMPPNP state that are coincident with enhanced dynamics in SecA itself (Fig. 5B.II). The decreased dynamics of the signal peptide throughout the ATPase cycle (Fig. 6E.I) are consistent with the client remaining largely tethered to the translocase via its signal peptide while mature domain parts associate/dissociate more dynamically (Burmann et al., 2013). Specifically, during the ATP hydrolysis transition (ADP:BeF_X_) state, all three regions of the client that bind SecA become more rigidified (Fig. 6EI-III), likely reflecting their being tightly caught in the ADP:BeF_X_-driven rigidified closed-flipped clamp (Fig. 5B.III). When ATP is hydrolyzed to ADP, as the translocase becomes more dynamic (Fig. 5B.IV), the grasp on the client chain relaxes modestly (Fig. 6E, brown to green).

In this recreated ATP hydrolysis cycle, each step of specific translocase conformations has a specific consequence on “catching and releasing” the three islands of the preprotein chain.

## Discussion

The gradual activation of the Sec translocase involves several interactions, including holo-enzyme assembly, nucleotide and client binding, dimer to monomer transitions that are orchestrated in hierarchical steps. Nonetheless, these steps can also occur in independent sub-reactions; e.g. clients can bind either onto quiescent-cytoplasmic or channel-primed SecA. How the Sec translocase and all complex nanomachines achieve hierarchical activation triggered promiscuously by hundreds of client interactions remains elusive. We reveal here a sophisticated two-part mechanism whereby, first various interactions triggered by clients work in concert to activate conformational switches within the translocase, and second those events lead to dynamics changes that ensure the translocation of the client.

Considerable evolutionary effort has prevented acquisition of a readily activated state in SecA_2_, favoring quiescence instead; e.g. by biased sampling of the wide-open clamp state (Krishnamurthy et al., 2021). Translocase activation relies on regulating pre-existing intrinsic dynamics of subunits that exist in a conformational ensemble (Ahdash et al., 2019; Allen et al., 2016; Corey et al., 2019; Gouridis et al., 2009). A full compendium of pre-existing conformations, arise from thermal atomic vibrations (Fig. 6C; S7C)(Bahar et al., 2010; Chen and Komives, 2021; Dobbins et al., 2008; Smit et al., 2021a; Smit et al., 2021b) and can be differentially sampled with minor energetic demands (Henzler-Wildman and Kern, 2007). Attesting to this, point mutations can recapitulate the effects of secretory preprotein binding to the translocase; e.g. trigger the translocase, stimulate its ATPase activity, drive monomerization (Gouridis et al., 2009; Gouridis et al., 2013; Karamanou et al., 2007). Prl mutations amongst them (Bieker et al., 1990; Bost and Belin, 1997; Schatz and Beckwith, 1990), shift the conformational landscape equilibrium of the holo-enzyme (Gouridis et al., 2009). For example, Prl mutations at gate2 of SecA or affecting the latter from a distance, mimic both the signal peptide/mature domain effects and facilitate ADP release (Fig. 2C).

At its core SecA can be seen as bearing two distinct modules, an ATPase hardwired onto a preprotein clamp (Fig. 7.I, grey) that assembles onto the SecY channel (yellow). Both the ATPase and the preprotein clamp display distinct local and domain dynamics, each finely controlled by different ligands (nucleotide or signal peptides), but these dynamics processes are largely uncoupled from each other. Only the arrival of non-folded, clients that bind to multiple patches on SecA couple these dynamics modules by providing a physical bridge between the ATPase and the clamp and by modulating them (Fig. 7.II). This coupling is controlled by dynamic switches (i.e. gate2 and Stem) that regulate the transduction of conformational signals downstream to effectuate the enzymatic activation, with the initiating step being ADP release (Fig. 7.III-IV), followed by the arrival of fresh ATP (Fig. 7.V). ATP cycle steps inside the motor have immediate ramifications on the conformational status of frustrated prongs in the clamp. These changes take the prongs through catch and release rounds of the client chain, at 2-3 distinct locations, thus biasing its forward motion (Fig. 7.III-VI).

**Figure 7.**
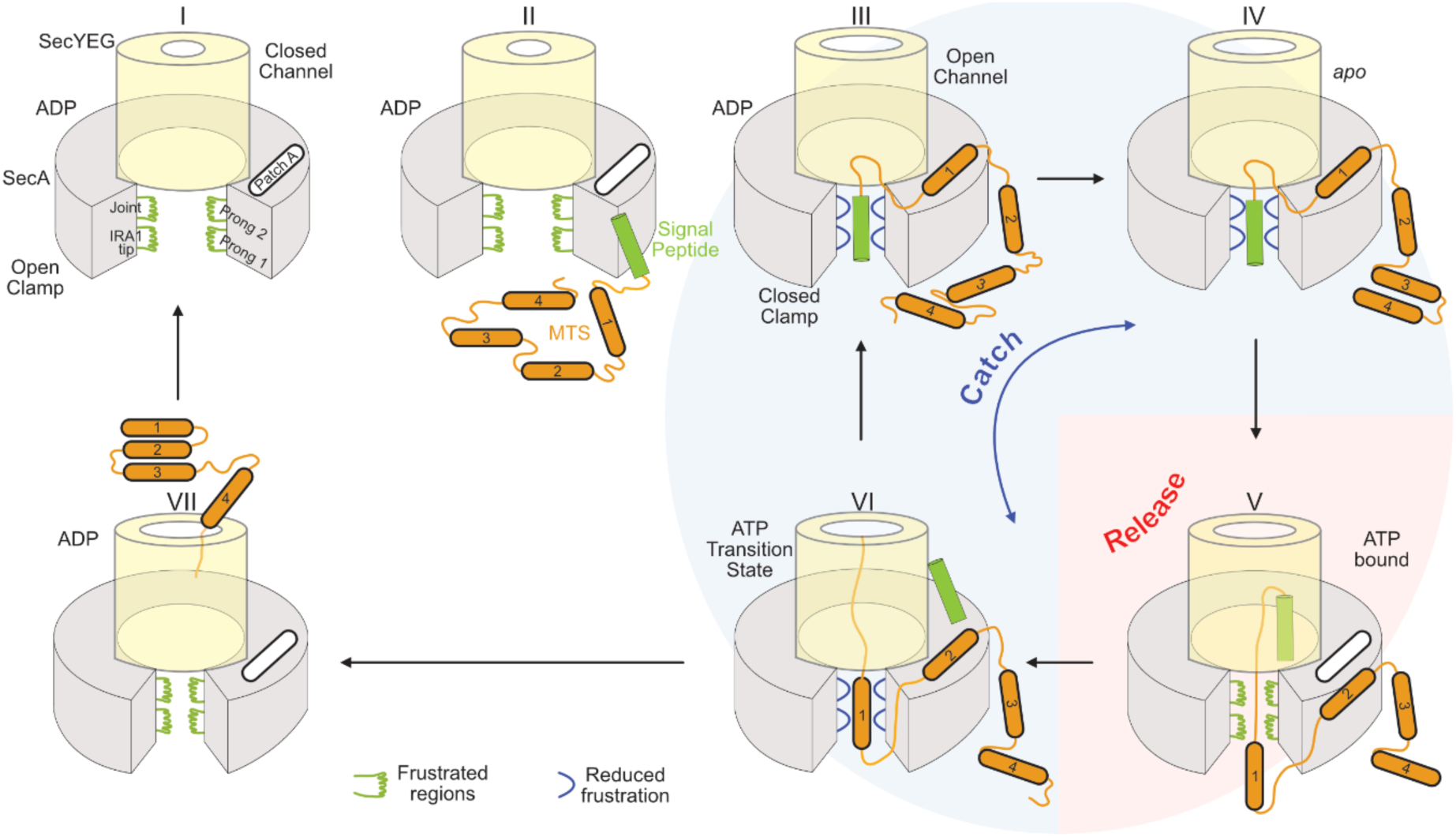
Dynamic regions on the translocase and model for preprotein translocation. Model of preprotein translocation by the Sec translocase. The primed translocase is in an open SecA clamp/closed channel state with frustrated client binding regions (green) (I). Preprotein binds onto the primed translocase (II). Signal peptide binding induces clamp closing and channel opening while MTS binding at PatchA (III) causes ADP release (IV). Binding is primarily through frustrated regions and hydrophobic interactions (PatchA). Client binding causes reduction in frustration (blue). ATP binding causes release of the chain and transport of the signal peptide into the channel (V), while MTS1 moves into the clamp and MTS2 binds to PatchA (VI). Processive ATP hydrolysis results in a series of ‘catch and release’ cycles, whereby MTSs serve as binding junctures on the translocase causing ADP release. The translocating chain occupies the channel and keeps it open during open/closed clamp states. Once the last MTS exits the channel, ADP cannot be released the translocase reverts back to a primed but dormant state (VII), awaiting the next client

Despite overall similarities between all polypeptides, only secretory preproteins are legitimate clients of the translocase in the cell. Their two moieties alone, signal peptides and mature domains, alter distinctly but only partially the conformational switches, thereby yielding inadequate functional effects (Fig. 3-4). Thus, isolated signal peptides shift the clamp towards closed states (Fig. 3A, lanes 10-12), partially elevate dynamics in gate2 (Fig. 2A-B) and loosen the channel (Fig. 7.II-III) (Duong and Wickner, 1999; Fessl et al., 2018; Gouridis et al., 2009; Knyazev et al., 2014). Mature domains alone increase local dynamics in the motor (Fig. S4G) and drive inefficient ADP release (Fig. 4D). However, only the synergy between the two covalently connected moieties, can efficiently couple the sub-reactions required to increase motor dynamics that facilitate efficient ADP release (Fig. 1B-C; C.II-III), a key rate limiting step (Fak et al., 2004; Robson et al., 2009; Sianidis et al., 2001). This mechanism ensures that random cytoplasmic proteins are not secreted.

ATP binding to the motor initiates translocation (Fig. 7.V). As the translocase cycles through nucleotide states, it manipulates the dynamics of its clients. In the ATP-bound state, the translocase retains signal peptide binding, releasing mature domain segments (Fig. 6E, red). In the transition state, it ‘catches’ both signal peptide and MTS regions tightly (Fig. 5B.III; 6B; 6E). Upon conversion to the ADP-bound state, a succeeding region of the bound client will induce ADP release (Fig. 1B). A new round of the nucleotide cycle starts. For as long as there are available succeeding mature domain segments to bind to the translocase and drive ADP release, the ATP hydrolysis cycle is repeated (Fig. 7.III-VI). In the absence of such segments, SecA remains in the quiescent ADP state, diffuses to the cytoplasmic pool and dimerizes (Gouridis et al., 2013). Preprotein translocation is achieved through such repetitive client-catch/release cycles regulated by nucleotide turnovers (Fig. 7). This mechanism is generic for both initiation and subsequent processive translocation steps (Fig. 1A.V). Only that in later stages of translocation, after signal peptide removal, they are simply replaced by internal hydrophobic MTS sequences (Chatzi et al., 2017)(Fig. 7.VI).

Our observations on the remarkable sensitivity of SecA dynamics to respond to even the slightest chemical change in the nucleotide invite a major rethink of the anticipated role of ATP hydrolysis in translocation. Rather than driving major deterministic strokes, a main role of nucleotides is revealed here to be the subtle step-wise regulation of the intrinsic dynamics of SecA (Fig. 5), by affecting intrinsically dynamic residues that line the nucleotide cleft (Keramisanou et al., 2006). Conformational cycles are regulated by the limited and transient interaction of the *γ*-phosphate of ATP and its transition states with NBD2 (Keramisanou et al., 2006; Papanikolau et al., 2007; Sianidis et al., 2001) and “locked” in its absence (Krishnamurthy et al., 2021), awaiting fresh client binding (Fig. 1B; Fig. 7.VI to I).

An under-appreciated property of secretory clients that enables them to modulate translocase properties, is their own elevated intrinsic dynamics. Mere reduction of these dynamics, in an otherwise binding-competent client, abrogated secretion (Sardis et al., 2017). We now reveal for the first-time islands in a secretory chain whose dynamics are differentially regulated in response to transient nucleotide states of the translocase (Fig. 6E). These segments encompass the signal peptide, MTSs and the region extending from the signal peptide C-region to the early mature domain [rheostat; (Sardis et al., 2017); Fig. 6D.I; 7.II]. Their detection was possible only because of the power of HDX-MS to analyze large membrane protein complexes in near-physiological conditions (Kochert et al., 2018) and derive medium resolution structural information on non-folded clients in translocating conditions (Tsirigotaki et al., 2017b), in the absence of detergents.

How does SecA promiscuously recognize/translocate its clients? Chaperones have been proposed to recognize frustrated regions in clients (He and Hiller, 2019; Hiller, 2019). Yet, frustrated regions also exist on chaperones; e.g. trigger factor (Ferreiro et al., 2014; Morgado et al., 2017) and identified here in SecA (Fig. 6B). The latter exhibits weak cytoplasmic/ATP-independent and strong membrane-associated/ATP-regulated holdase activity (Gouridis et al., 2009). Frustrated regions, of both clients and chaperones, can sample a wide conformational and sequence space until they interact tightly (Ferreiro et al., 2014; He and Hiller, 2018). Thus, a chaperone can promiscuously interact with hundreds of clients without a need for rigid lock and key recognition. Such a mechanism is optimally suited to dealing with non-folded clients. In the case of SecA, the three highly dynamic, locally frustrated prongs on either side of the clamp and the tip of IRA1 (Fig. 6A-B; Fig. 7.I-III), the electronegative environment of the clamp (Fig. S5B) and the adjustability of its width due to PBD and NBD2 rigid body motions, enhance plasticity and possible interactions with non-folded clients, potentially enabling accommodation of even partially locally folded structures (Tsirigotaki et al., 2018). Client-translocase interactions reduce dynamics both in the frustrated prongs of SecA (Fig. 5B.III; Fig. 7, green) and the transiently structured/frustrated elements of the client (Fig. 6D). A powerful aspect of this mechanism is that interaction with the client can be of high affinity yet transient, quickly returning to a looser “release” state. We hypothesize that the combination of multiple frustrated prongs and multiple recognition patches on the client, form an optimal basis of SecA processivity, that would have otherwise been impeded by tighter/extensive recognition solutions of an unfolded polymer.

SecA acts as a mechanical device that biases vectorial forward motion. This is not common among soluble chaperones. Presumably, local interactions of frustrated clamp regions are sufficient to stall backward sliding of the exported chain yet loose enough to allow forward motion of untethered segments through the channel. This ‘catch and release’ mechanism is an important feature of the translocation process. Release cycles allow chain segments to enter the channel; catch cycles would bind a downstream segment and prevent back-slippage (Fig. 7.V and VI). This mechanism is compatible either with a “brake” preventing backsliding (Vandenberk et al., 2019) and allowing Brownian “ratchet” motion of the unbound parts (Allen et al., 2016; Allen et al., 2020) or as part of a power stroke, if catching actively carries along chain segments further into the channel (Catipovic et al., 2019). This mechanism is also compatible with a continuum ratchet model, where the SecA motor moves stochastically along a periodic potential, coupled to the ATP cycling, providing the required time correlation necessary for net vectorial motion (Magnasco, 1993). All these models would presumably include MTSs (Chatzi et al., 2017) and possibly other “catch” signals (Fig. 6D-E).

We focused here on a short preprotein that cannot fold extensively. This allowed us to dissect translocase binding away from the interference of folding propensities (Tsirigotaki et al., 2018). It is anticipated that the same fundamental mechanisms apply in steps of longer mature domain translocation. The mere presence of the mature chain trapped inside the channel effectively forces the channel to maintain a “loose” state (Fig. 7.V), even in the later stages of translocation after the signal peptide has been cleaved/ejected (Schiebel et al., 1991). Chaperones bind to fast folding clients and regulate their conformational collapse (He et al., 2016; Kellner et al., 2014). Secretory proteins on the other hand are highly flexible with long-lived folding intermediates (Tsirigotaki et al., 2018). Nevertheless, for clients that fold rapidly or have domains that might fold while their N-terminus is being translocated, translocase dynamics may serve to counter inherent folding forces in the client (Arkowitz et al., 1993; Gupta et al., 2020) alone, or in concert with other chaperones (De Geyter et al., 2020; Fekkes et al., 1997).

## Experimental Procedures

### Materials

For buffers, strains, plasmids and primers see Supplementary Table S3, S4, S5 and S6 respectively. TCEP ([Tris(2-carboxyethyl)phosphine] was from Carl Roth, formic acid-MS grade from Sigma Aldrich, LC-MS grade acetonitrile LC-MS grade from Merck. D_2_O (99.9%) was obtained from Euroisotop; Alexa555 and Alexa647 - maleimide from Thermo Fisher Scientific. All other chemicals and buffers were ACS grade from Merck or Carl Roth. proPhoA signal peptide, obtained from GenScript as lyophilized powder, was dissolved in DMSO (Merck) to a final concentration of 45 mM. Protein concentration was determined using either Bradford assay (Biorad) or/and Nanodrop^TM^ spectrophotometry (Thermo Scientific) following manufacturer’s instructions. MANT-ADP was from Invitrogen/Thermo Fisher Scientific.

### Molecular cloning

Site directed mutagenesis was performed using QuickChange site directed Mutagenesis protocol (Stratagene Agilent) using indicated vector templates and primers. Molecular cloning and sample handling was as previously described (Krishnamurthy et al., 2021)

### Protein purification

SecA and derivatives were overexpressed in T7 Express lysY/I^q^ [derivative of BL21 (DE3)] cells and purified as described (Papanikolau et al., 2007). All proteins were assessed for purity and quality using gel filtration chromatography and SDS-PAGE. The His-tagged derivatives of SecA-D2, proPhoA_1-122_ and PhoA_23-122_ were purified as previously described (Chatzi et al., 2017; Vandenberk et al., 2019). His-SecD2 were stored as described (Krishnamurthy et al., 2021), while proPhoA_1-122_ and PhoA_23-122_ were stored in buffer A (Chatzi et al., 2017).

SecYEG-IMVs and derivatives were prepared as in (Lill et al., 1989; Lill et al., 1990) and concentration was determined as described (Gouridis et al., 2013). All biochemicals were tested for functional activity in ATPase and *in vitro* preprotein translocation assays.

### MANT-ADP release assays

MANT-ADP release assays were carried out as described in (Krishnamurthy et al., 2021). SecYEG:SecA_2_ (0.5 µM) were added to MANT-ADP (1 µM; 30 s) to initiate MANT-ADP binding onto the translocase. Client proteins were added (chase; 90 s) at the following final concentrations: proPhoA_1-122_ – 15 µM; PhoA_23-122_ – 20 µM; signal peptide – 30 µM.

### Dynamics of the Sec translocase by HDX-MS

HDX-MS experiments were carried out as previously described (Krishnamurthy et al., 2021). SecA, all their derivatives were diluted into buffer B to a final concentration of ∼100 µM prior to HDX-MS analysis. To monitor SecA:proPhoA_1-122_ interactions in solution, proPhoA_1-122_ (in Buffer A) was diluted in buffer B to a final concentration of 250 µM (0.2 M Urea), immediately added to SecA at 4 µM: 35 µM ratio (SecA: proPhoA_1-122_) and incubated for 2 minutes prior to D exchange. Complexes of the channel with SecA and its derivatives were generated and analyzed as described (Krishnamurthy et al., 2021). To monitor how signal peptides (SP) activate the translocase, the synthetic proPhoA SP (Genescript; 45mM in 100 % DMSO) was diluted 30-fold into Buffer B (to obtain 1.5 mM SP in 3 % DMSO), added to preincubated SecA:SecYEG at a final molar ratio of 4 µM: 6 µM: 30 µM (SecA:SecYEG:SP) and the reaction was incubated for a further 1 minute.

SecYEG:SecA:client interactions: To monitor the dynamics of SecA as part of SecYEG:SecA:client complexes, the client (proPhoA_1-122_ and PhoA_23-122_) were added in excess to preincubated SecYEG:SecA to maintain a final molar ratio of 4 µM: 6 µM: 20 µM. In SecYEG:SecA: signal peptide + mature domain complexes, proPhoA signal peptide was added to preincubated SecYEG:SecA:PhoA_23-122_ (as described above) to a final concentration of 30 µM. Indicated concentrations are in the final D-exchange reaction. D-exchange labeling was carried out in D_2_O buffer C at 30 °C for 7 timepoints (10 s, 30 s, 1 min, 2 min, 5 min, 10 min, 30 min) in 3 technical replicates (Table S1). Reaction was quenched in buffer D and HDX-MS data acquisition and analysis was performed as described (Krishnamurthy et al., 2021). All mutant proteins were handled similar to the wild type ones and reactions were maintained at similar molar ratios.

### Dynamics of client proteins by HDX-MS

To monitor dynamics of free proPhoA_1-122_ and PhoA_23-122_, proteins were diluted from 6 M urea into buffer B to a final concentration of 50 µM, and subsequently diluted 10-fold into D_2_O buffer E. D-labeling was carried out for a short 10 s pulse at 4 °C.

To monitor dynamics of client proteins when bound to the Sec translocase (see Fig. S7A for experimental schematic), to ensure all available client proteins are bound to translocase, the concentration of client proteins was maintained sub stoichiometric to the translocase. Prior to D-exchange, the complete SecYEG:SecA;preprotein complex is generated by incubating SecA_2_ (20 µM) with SecYEG (40 µM) for 5 min on ice. Client proteins were added to a final concentration of 15 µM and incubated for 5 min on ice.

The complex was incubated for 20 s at 37 °C (this step was omitted for low temperature experiments). D-exchange reaction was initiated by diluting the complex 10-fold in D_2_O buffer C. Labeling was carried out for 10 s at 4 °C. Reaction was quenched using buffer D, proteins were proteolyzed and injected into a HDX sample manager (Waters, Milford) for UPLC based peptide separation as described (Krishnamurthy et al., 2021). To unambiguously detect the low abundance peptides from the client proteins, data acquisition was carried out in UDMS^E^ data acquisition mode. Data acquisition parameters were as described (Cryar et al., 2017). Peptide analysis and quantification was as described (Krishnamurthy et al., 2021).

### Determination of ΔG_ex_ values

ΔG_ex_ values were determined using PyHDX software (v0.4.0-rc1) (Smit et al., 2021a). D-uptake data from triplicate experiments, for all timepoints were input along with 100% deuteration control. Input parameters were set to the following parameters: temperature - 303 K; ph – 8; stop loss – 0.01; stop patience – 100; learning rate – 10; momentum – 0.5; epochs – 100000; regularizer 1– 0.01, regularizer 2 – 0.01.

### Single-molecule fluorescence microscopy and PIE

Protein purification, fluorescent labeling, sample preparation and data analysis, quantification and statistical analysis for single molecule PIE based FRET experiments were carried out as previously described (Krishnamurthy et al., 2021). Confocal scanning analysis mode was applied to follow the conformational dynamics of SecA in solution at 20°C. To follow the effect of signal peptide to clamp dynamics of the translocase, signal peptide was added to monomeric, dimeric or channel-primed SecA [generate as described (Krishnamurthy et al., 2021)] to a final concentration of 37 µM. proPhoA_1-122_ was added to a final concentration of 10 µM. All mutant derivatives were handled similar to wild-type proteins.

### H-bonding graph analysis

To determine H-bond paths and long-distance conformational couplings between the signal peptide binding cleft and gate2, we used algorithms based on graph theory and centrality measures as described (Karathanou and Bondar, 2019; Krishnamurthy et al., 2021). Briefly, residues were considered H-bonded if the distance between the hydrogen and acceptor heavy atom, d_HA_, is ≤2.5 Å. H bonds were calculated between protein sidechains, and between backbone groups and protein sidechains. Data are visualized as H-bond networks with unique lines (coloured according to H-bond frequency) between Cα atoms of residue pairs that H-bond. H-bond frequency is the percentage of analysed trajectory segment during which two residues are H-bonded.

### Normal Mode Analysis

Normal modes describe protein vibrational movements, were calculated using the WebNM@ web server (Tiwari et al., 2014) with PDB ID: 2VDA as the input structural model if SecA. Per-residue displacement and normal mode flexibility were derived from normal mode eigenvalues as described (Dobbins et al., 2008; Smit et al., 2021a). Total vibrational displacement of each residue undergoing fluctuations under low frequency normal modes (modes 7-12) are calculated and plotted. Residues that undergo displacement greater than 2 are highlighted in shades of blue.

### Miscellaneous

Pymol (https://pymol.org/) was used for structural analysis and visualization. SecA activation energy determination, *in vivo* proPhoA and PhoA translocation, *in vitro* proPhoA translocation, SecA ATPase activity, *in vivo* SecA complementation, affinity determination of SecA and/or proPhoA for the translocase, were as described (Chatzi et al., 2011; Gouridis et al., 2009; Gouridis et al., 2010; Gouridis et al., 2013). H-bond networks between the signal peptide cleft and motif IVa were determined as described (Krishnamurthy et al., 2021). ADP:BeF_X_ was generated by adding ADP:BeCl_2_:NaF in a 1:1:5 ratio and incubated at 4 °C for 30 min to obtain a final concentration of 50 mM.

## Acknowledgements

We are grateful to: T. Cordes for sharing software for smFRET data analysis. Our research was funded by grants (to AE): MeNaGe (RUN #RUN/16/001; KU Leuven); ProFlow (FWO/F.R.S.-FNRS “Excellence of Science - EOS“ programme grant #30550343); DIP-BiD (#AKUL/15/40 - G0H2116N; Hercules/FWO); CARBS (#G0C6814N; FWO); Profound (WoG Research Training Network, FWO, Protein folding/non-folding and dynamics; #W002421N) and (to AE and SK): FOscil (ZKD4582-C16/18/008; KU Leuven) and (to A-NB): by the Excellence Initiative of the German Federal and State Governments via the Freie Universität Berlin, and by allocations of computing time from the North-German Supercomputing Alliance, HLRN. SKr was a FWO [PEGASUS]² MSC fellow; NE was a MSCA SoE FWO fellow; JHS is a PDM/KU Leuven fellow; GG was a Rega Foundation postdoctoral program fellow. This project has received funding from the Research Foundation – Flanders (FWO) and the European Union’s Horizon 2020 research and innovation programme under the Marie Skłodowska-Curie grant agreements No 665501 and 195872.

## Author contributions

SKr purified proteins and membranes, did biochemical and fluorescence assays, designed and performed HDX-MS work and data analysis. MFS and KEC purified proteins, performed molecular biology, *in vivo* and *in vitro* biochemical and biophysical assays. NE purified and labelled proteins and performed smFRET experiments and data analysis. JHS developed PyHDX software and analysed HDX-MS data, adapted FRET burst analysis for Microtime200 output data and performed NMA analysis. KK performed MD simulations and graph analysis of H-bond networks. GG performed biochemical, molecular biology and biophysical assays, analysed data and advised on smFRET. AGP performed molecular cloning and mutagenesis. ANB set up and supervised the MD simulations and graph analysis. SK designed and supervised molecular biology experiments, purified proteins, performed biochemical and biophysical assays and data analysis. AE did structure and data analysis and designed experiments. SKr and AE wrote the first draft and finalized it with contributions from SK, ANB, JHS and NE. All authors reviewed and approved the final manuscript. AE and SK conceived and managed the project.

