## Supplementary material for "Preproteins couple the intrinsic dynamics of SecA to its ATPase cycle to translocate via a catch and release mechanism"

**Running title:** Preprotein-coupled translocase dynamics

**Keywords:** translocase; protein secretion; SecA; SecYEG channel, HDX-MS, smFRET, Molecular Dynamics, signal peptide, secretory clients, intrinsic dynamics, enzyme activation

**Table of contents**

**Supplemental figures**……………………………………………………………………p.3
**Figure S1:** Preproteins induce ADP release by allosterically enhancing the dynamics of the ATPase motor (related to Fig. 1)
**Figure S2:** Signal peptides trigger the translocase by enhancing gate2 dynamics (related to Fig. 2)
**Figure S3:** Clamp closing is a major conformational event towards translocase triggering (related to Fig. 3)
**Figure S4:** Fig. S4: Mature domain-driven ADP release and ATP turnovers (related to Fig. 3 and 4)
**Figure S5:** Nucleotide occupancy and preproteins drive translocase conformational motions (related to Fig. 5)
**Figure S6:** Nucleotide regulated dynamics in channel-bound SecA and structure of the closed flipped clamp state (related to Fig. 5)
**Figure S7:** Translocase binds and regulates preprotein dynamics (related to Fig. 6)

**Supplemental tables**……………………………………………………………………p.17
**Table S1** Dynamics of the Sec translocase by HDX-MS

**Table S2** Dynamics of client proteins by HDX-MS

**Table S3** List of buffers

**Table S4** List of strains

**Table S5** List of plasmids

**Table S6** List of primers

**Supplementary Movies**
**Movie S1** Clamp motions of the translocase (related to Fig. 3 and S6)

#### Supplementary Figures

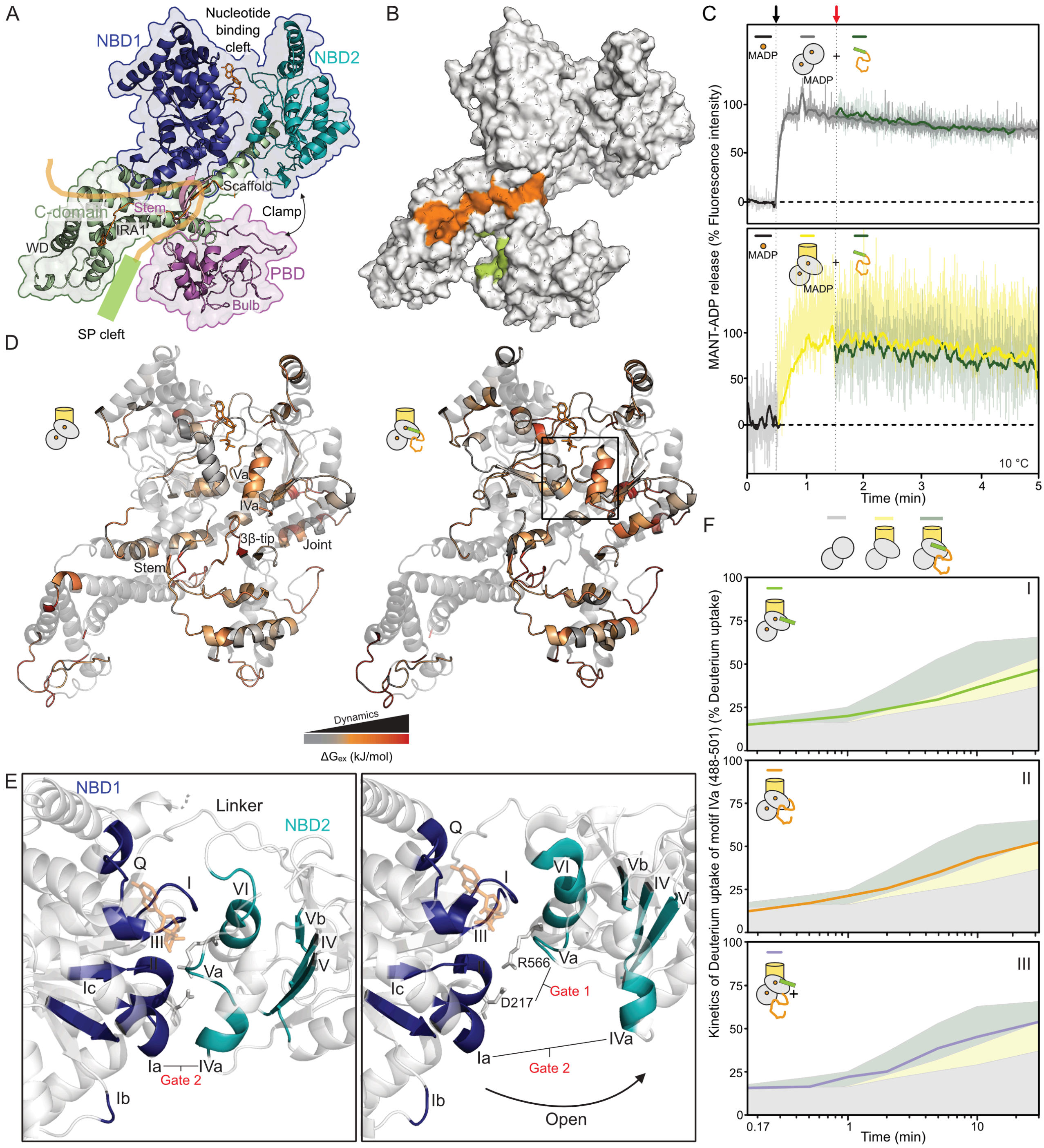


### Figure S1 Preproteins induce ADP release by allosterically enhancing the dynamics of the ATPase motor (related to Fig. 1)

**A.** Domain organization of *ec*SecA (PDB: 2VDA; the C-tail was obtained from 1M6N), shown in ribbon representation. NBD: Nucleotide binding domain 1 and 2 are Rossman fold domains that together form the ATPase motor that is homologous to that of DEAD-box helicases particularly those of superfamily II (blue outline) (Papanikou et al., 2007); PBD: preprotein binding domain. The regulatory C-domain comprises the Scaffold (SD), Wing (WD), IRA1 (Intra-molecular regulator of ATPase 1) and C-tail. The C-tail loops back to form a β-sheet with the Stem. ADP is shown as orange sticks. A bound preprotein is shown; signal peptide (green), mature domain (orange).

**B.** Preprotein binding sites, mapped onto the space-filling representation of SecA. The signal peptide binding site in the PBD bulb is green (Gelis et al., 2007). The mature domain binding onto PatchA, a hydrophobic surface spanning from Stem to IRA1 and WD (Chatzi et al., 2017), is orange.

**C.** ADP release assays follow the fluorescence intensity of MANT-ADP upon binding to SecA (as in Fig. 1B).
Top: $SecA_2$ (0.5 μM) addition (black arrow at 30 s) to free MANT-ADP (1 μM), at 37 °C results in an increase in fluorescence intensity. Addition of $proPhoA_{1\text{-}122}$ (15 μM; green line) chase (red arrow at 90 s) does not change the fluorescence intensity, suggesting that binding of preprotein to soluble SecA does not cause ADP release.
Bottom: Channel bound $SecA_2$ binds to MANT-ADP at 10 °C and increases its fluorescence intensity (yellow line; as in Fig.1B). However, at 10 °C, addition of $proPhoA_{1\text{-}122}$ (15 μM; green line) chase (red arrow at 90 s) does not change the fluorescence intensity, unlike what was observed at 37°C (Fig.1B). Clearly, preprotein binding does not cause ADP release from channel-primed $SecA_2$ at non-physiological temperature.

**D.** Dynamics of SecYEG:$SecA_2$:ADP (left) and SecYEG:$SecA_2$:ADP:$proPhoA_{1\text{-}122}$ (right). $\Delta G_{ex}$ values, calculated per residue using the PyHDX software (Smit et al., 2021) (as in Fig. 1D), were colour-mapped onto the Closed clamp structure of SecA (PDB: 3DIN). Rigid regions (grey; $\Delta G_{ex}$ ~29 kJ/mol), flexible (orange; $\Delta G_{ex}$ ~21 kJ/mol), dynamic (red; $\Delta G_{ex}$ ~12 kJ/mol). Boxed area: gate2.

**E.** Helicase motifs onto the ATPase motor of SecA are colour indicated; motifs in NBD1 (I, Ia, Ib, Ic, II, III) are in blue, those in NBD2 (IV, IVa, V, Va, Vb, VI) in teal. The flexible linker that connects NBD1 and 2 is indicated. The motor predominantly exists in a closed form (left). NBD2 swings outwards resulting in an open motor (PDB: 2FSI). Gate1: salt bridge between D217 and R566, regulates the ATPase activity of SecA (Karamanou et al., 2007), Gate2: a contact between motif $Ia_{NBD1}$ and motif $IVa_{NBD2}$ (discussed below). Disrupting the Gate2 interaction results in motor loosening, to an 'open' state.

**F.** D-uptake kinetic plots of a motif IVa peptide (aa488-501), shown as a percentage of its full deuteration control (Table S1) (as in Fig. 1E). Addition of signal peptide (I) or mature domain (II) alone, or in combination *in trans* (III), to the SecYEG:$SeA_2$:ADP state marginally alter the dynamics of motif IVa peptide (compare yellow to line). The indicated states are shown on top of a transparent Fig. 1E acting as reference. SD error values are within the width of the data point and hence are not visible. *n = 3*.

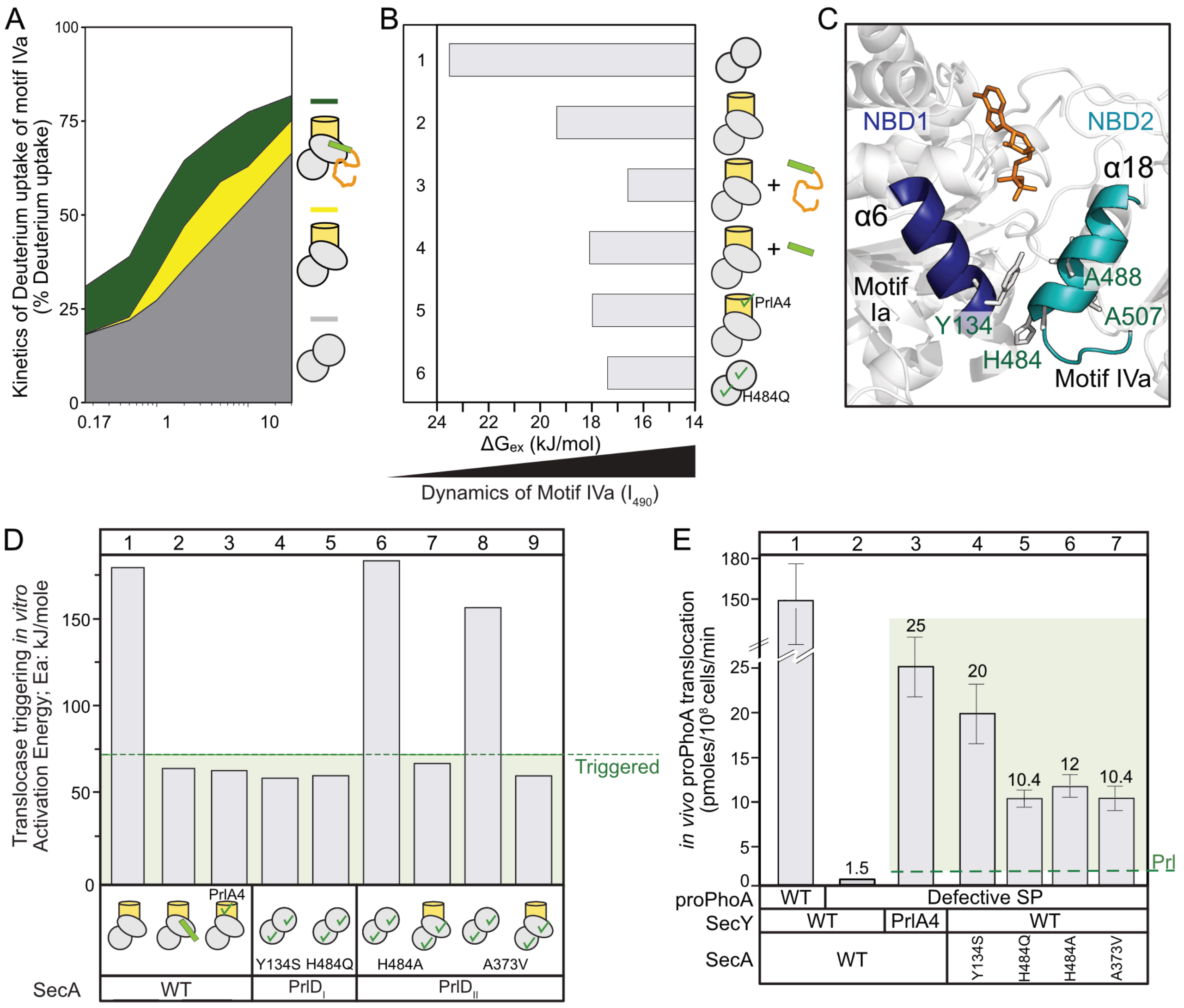


**Figure S2 Signal peptides trigger the translocase by enhancing gate2 dynamics (related to Fig. 2)**

**A.** D-uptake kinetic plots of a motif IVa peptide (aa 488-501), shown as a percentage of its full deuteration control (Table S1), for $SecA_2$ (grey), $SecYEG:SecA_2$ (yellow) and $SecYEG:SecA_2$:preprotein (dark green), in the absence of ADP. The kinetics of the peptide retain the trends seen in Fig. 1E upon addition of channel and preprotein, with minor differences between the ADP presence/absence.

**B.** $\Delta G_{ex}$ values were calculated for residue $I_{490}$ of motif IVa for the indicated conditions (as in Fig. 1F), in the absence of ADP. The lower the $\Delta G_{ex}$ values the more increased the dynamics of $I_{490}$. Channel binding to $SecA_2$ results in 4 kJ/mol reduction (compare lane 1 and 2). Preprotein binding to $SecYEG:SecA_2$ results in a further 2 kJ/mol reduction (lane 3) while signal peptide binding only in a 1 kJ/mol further reduction (lane 4). $SecY_{PrlA4}EG:SecA_2$ (lane 5) and soluble $SecA(H484Q)_2$ (lane 6) exhibit $\Delta G_{ex}$ values as low as the $SecYEG:SecA_2$:signal peptide.

**C.** Motif Ia on α6 of NBD1(dark blue) and motif IVa on α18 of NBD2 (cyan) come together to form Gate2. More specifically, the latter is formed by an interaction between $Y134_{NBD1}$ with $H484_{NBD2}$ and $A488_{NBD2}$. All three residues are known PrlD mutants (in green; protein localization mutants in SecA; (Huie and Silhavy, 1995); A507 another known PrlD residue lies close to the Gate2 contact. ADP is shown as orange sticks.

**D.** Activation Energies ($E_a$) of wild type, or the indicated mutant, translocases under translocation conditions (0.4 μM SecA; 1 μM SecYEG supplemented with 9 μM proPhoA signal peptide, (indicated by green tube) derived from Arrhenius plots, as described (Gouridis et al., 2009). *n=3*.

**E.** *In vivo* secretion of the signal peptide defective pro(L8Q)PhoA driven by wt (lane 2) or PrI translocases (lanes 3-7; as indicated). Alkaline phosphatase units were converted to protein mass, as described (Gouridis et al., 2010). pro(L8Q)PhoA is not secreted by the wild type translocase (lane 2) but is secreted by PrI translocases (lanes 3-7). Translocation of wt proPhoA by the wild type translocase under the same conditions (lane 1) is shown. *n*= 6; mean values (± SEM). PrI mutants translocate more than 3 pmoles/ $10^8$ cells/ min of defective pro(L8Q)PhoA (green shade).

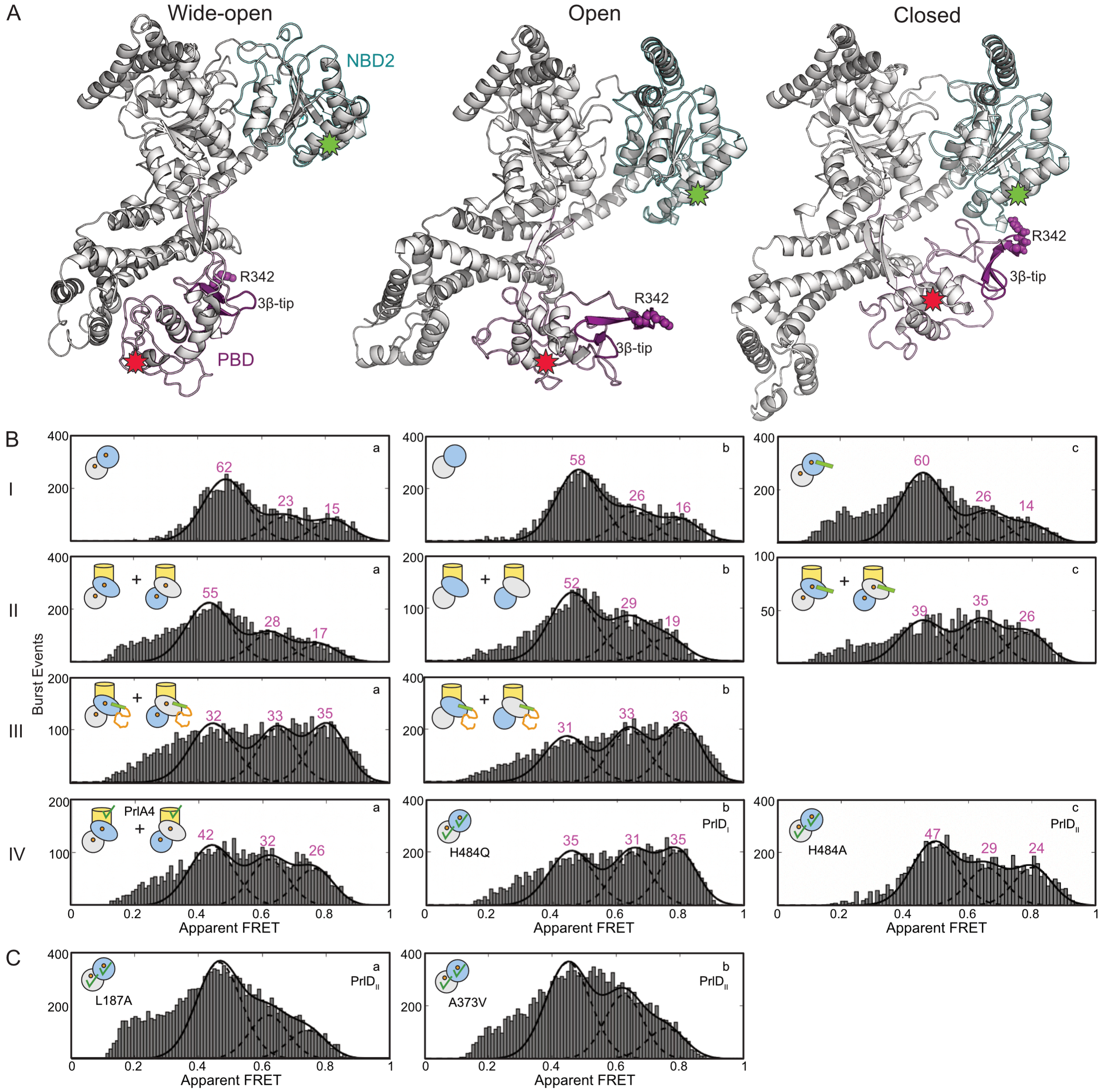


**Figure S3 Clamp closing is a major conformational event towards translocase triggering (related to Fig. 3)**

**A.** The protomer of *ec*SecA structure is used to visualize the three known clamp states: Wide-open (modelled from PDB: 1M6N), Open (PDB: 2VDA) and Closed (modelled from MD simulations; (Krishnamurthy et al., 2021). The position of fluorophores used in smFRET experiments is indicated with red (V280C) and green (L464C) stars. During labeling the acceptor/donor fluorophores can stochastically occupy either location; only molecules carrying both fluorophores were analyzed. R342 (magenta spheres) and the 3β-tip (magenta) in PBD, that move close to/interact with NBD2 as the clamp closes are indicated. PBD is outlined in magenta, NBD2 in cyan.

**B-C.** PIE-smFRET solution experiments of SecA under the indicated conditions (by pictograms) are presented as 2D plots. Histograms,apparent FRET value ($E^*$; x-axis) against burst events (y-axis), were fitted with Gaussian distributions as described (Krishnamurthy et al., 2021). The area under the curve was quantified and expressed as a percentage (pink numbers) of the sum of areas under all curves.

Soluble and channel primed SecA were generated as described (Krishnamurthy et al., 2021). Signal peptide (37 μM) or proPhoA$_{1-122}$ (10 μM), were incubated with free or channel bound SecA, wild type or mutant derivatives (as indicated; 5 min, 4 °C).

*Inlet pictograms indicate*: the labeled SecA protomer (blue) and under consideration (apoprotein or ADP- or signal peptide/preprotein- bound; freely diffusing or channel bound). Channel-bound SecA exists as two forms, the active protomer binds to SecYEG (oval) the other one (circle) is regulatory (Krishnamurthy et al., 2021). For analysis of mutants, SecA homodimers (wild type) are compared to heterodimers (the labelled SecA protomer is the indicated mutant while the non-labelled protomer is wild type). SecA: circles/ovals; SecYEG: yellow cylinder; Prl mutations: green ticks; ADP: orange circle; signal peptide: green cylinder; preprotein: green cylinder with orange line. Pink numbers: quantification (%) of SecA's clamp states

**B.** Clamp states of SecA monitored at the indicated regimes.

**C.** Clamp states of the indicated Stem/α8 interface mutants of SecA.

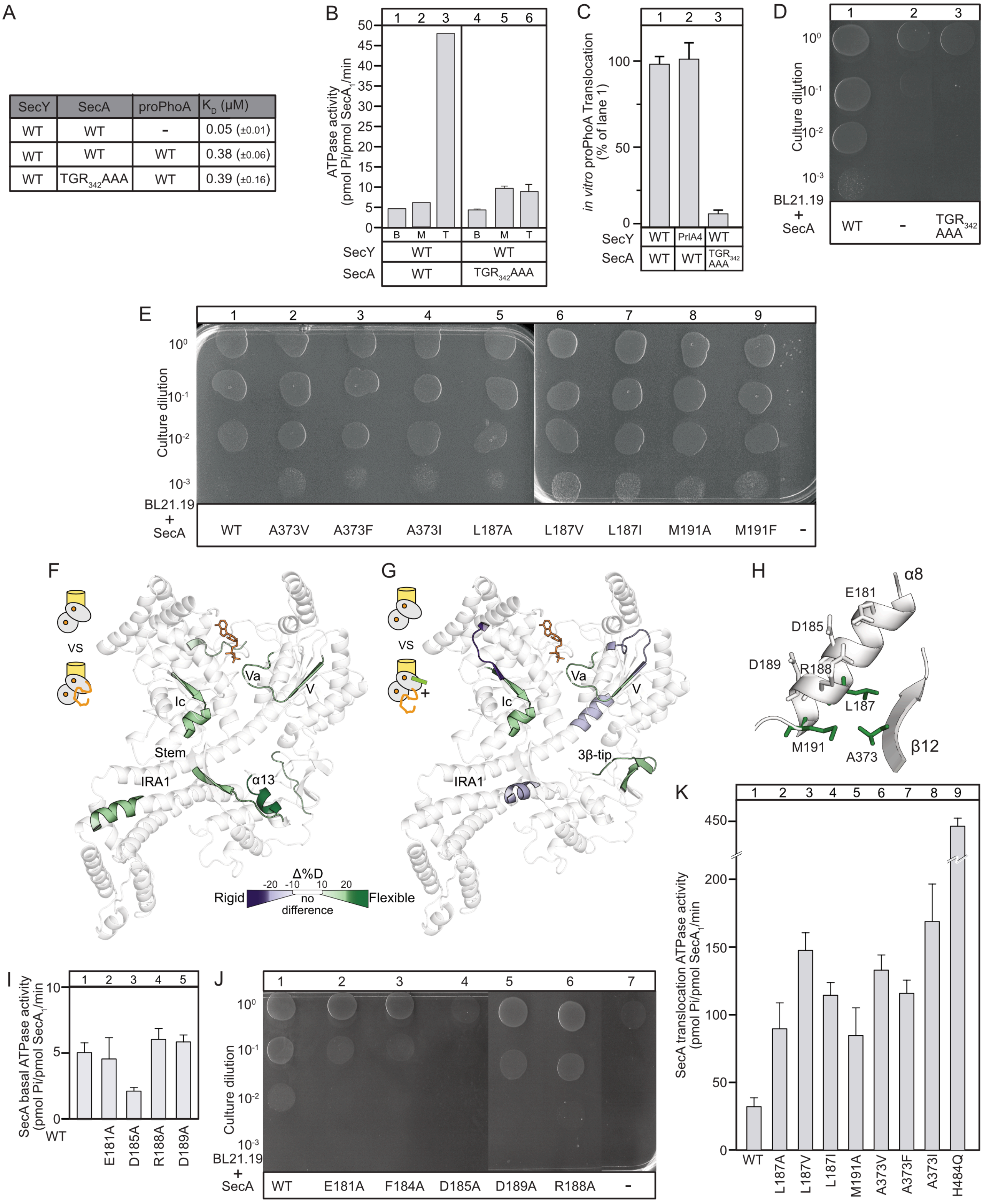


| SecY | SecA | proPhoA | $K_D$ (μM) |
|---|---|---|---|
| WT | WT | - | 0.05 (±0.01) |
| WT | WT | WT | 0.38 (±0.06) |
| WT | $TGR_{342}AAA$ | WT | 0.39 (±0.16) |

**Figure S4: Mature domain-driven ADP release and ATP turnovers (related to Fig. 3 and 4)**

**A.** Binding affinities ($K_D$) of SecA for the SecYEG channel and of channel-bound wild type SecA or SecA($TGR_{342}AAA$) for proPhoA. *n*= 6; mean values (± SEM).

**B.** The ATPase activity of the freely diffusing (basal; B; 0.4μM SecA), channel-bound (membrane; M; 1 μM SecY) and translocating (T; SecY plus 9 μM proPhoACys$^-$) wild type

SecA or SecA($TGR_{342}AAA$) was determined at 37 °C, as described (Gouridis et al., 2010). n= 6; mean values (± SEM).

**C.** *In vitro* translocation of proPhoACys$^-$ (9μM) driven by wild-type or mutant translocases (as indicated), at 37°C, as described (Gouridis et al., 2010). The polypeptide amount translocated by the wild-type translocase was considered 100%; all other values were expressed as a percentage of this. *n= 3*; mean values (± SEM).

**D-E.** *In vivo* genetic complementation of the *E.coli* BL21.19*secAts* strain by either an empty vector or one carrying *secA* wt or (**D**) SecA($TGR_{342}AAA$) or (**E**) Stem/α8 mutants, as indicated. Serial dilutions of the cultures ($OD_{600}$=0.5) were spotted (12μl) on LB-Ampicillin plates and grown at 42°C. *n*= 3; a representative experiment is shown. All spots shown are from the same biological replicates. Extraneous lanes have been removed resulting in image splicing lines.

**F.** Effect of PhoA(23-122) (15 μM) binding on the local dynamics of $SecA_2$:ADP:SecYEG (shown as in Fig. 1C).

**G.** Local dynamics effect of proPhoA signal peptide (30 μM) and PhoA(23-122) added *in trans* onto $SecA_2$:ADP:SecYEG (shown as in Fig. 1C)

**H.** Ribbon representation (PDB: 2VDA) of the Stem(β12)/α8 interface of SecA, with residues shown in stick representation. Residues that face the interface are in green; the rest in grey.

**I.** Basal ATPase activity of freely diffusing SecA and the indicated α8 mutants (as in B).

**J.** *In vivo* genetic complementation of the *E.coli* BL21.19*secAts* strain using wild type SecA or the indicated α8 mutants (as in C).

**K.** Translocation ATPase assay (as in B) for wild type SecA and the indicated Stem/α8 mutants showing preprotein hyper-stimulated ATPase activity.

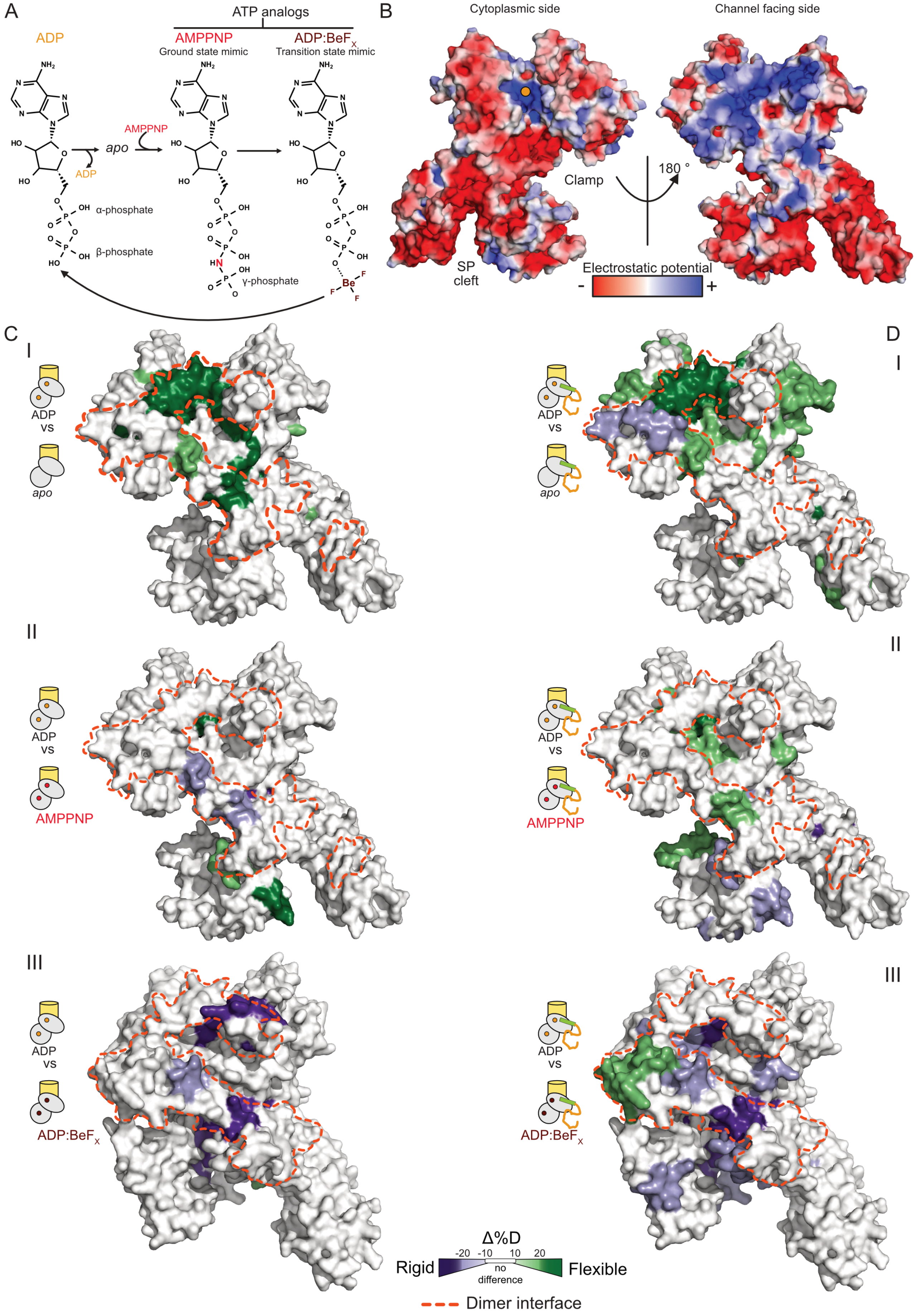
A
ATP analogs
ADP
AMPPNP
Ground state mimic
ADP:BeF$_X$
Transition state mimic
AMPPNP
apo
ADP
α-phosphate
β-phosphate
γ-phosphate
B
Cytoplasmic side
Channel facing side
Clamp
180 °
SP cleft
Electrostatic potential
-
+
C
D
I
ADP vs apo
II
ADP vs AMPPNP
III
ADP vs ADP:BeF$_X$
Δ%D
-20 -10 10 20
Rigid
no difference
Flexible
Dimer interface

**Figure S5 Nucleotide occupancy and preproteins drive translocase conformational motions (related to Fig. 5)**

**A.** The nucleotide cycle in the translocase is initiated by the release of ADP to yield the *apo* form. The ATP bound but not yet hydrolyzed state is mimicked by AMPPNP, a non-hydrolyzable analog of ATP with an N atom (red; middle panel) in place of O. ADP:$BeF_X$ is considered a pre-hydrolysis transition state analog with the $BeF_X$ moiety (brown) mimicking the γ-phosphate (Lacabanne et al., 2020). Hydrolysis of ATP results in the formation of ADP that needs to be released by the next segments of the preprotein clients, thereby restarting the nucleotide cycle. The key difference between the nucleotides lies in the presence/position of the γ-phosphate group that can interact with R509 in NBD2 (Papanikolau et al., 2007).

**B.** The electrostatic potential of SecA (PBD: 2VDA) was calculated by APBS electrostatics calculations (Baker et al., 2001) and mapped onto the structure. The cytoplasm-facing side of SecA when it is monomeric and bound to SecY (left) is predominantly negatively charged, with the signal peptide cleft and the inside of the clamp particularly negatively charged. The nucleotide binding cleft (left; orange circle) is positively charged. The membrane and SecY channel facing side of SecA (right; top) is predominantly positively charged.

**C-D.** The structures shown in Fig. 5A and 5B, are flipped 180 ° here, to show the dimerization interface (indicated by red dashed lines; derived from ecSecA modelled after PDB: 1M6N) (Gouridis et al., 2013; Krishnamurthy et al., 2021) drawn on the SecY-binding side (Zimmer et al., 2008) once SecA becomes monomeric. Nucleotides cause varying dynamics responses at the dimerization interface, that appear localized in regions without affecting the entire interface, indicating quaternary dynamics effects between the two protomers.

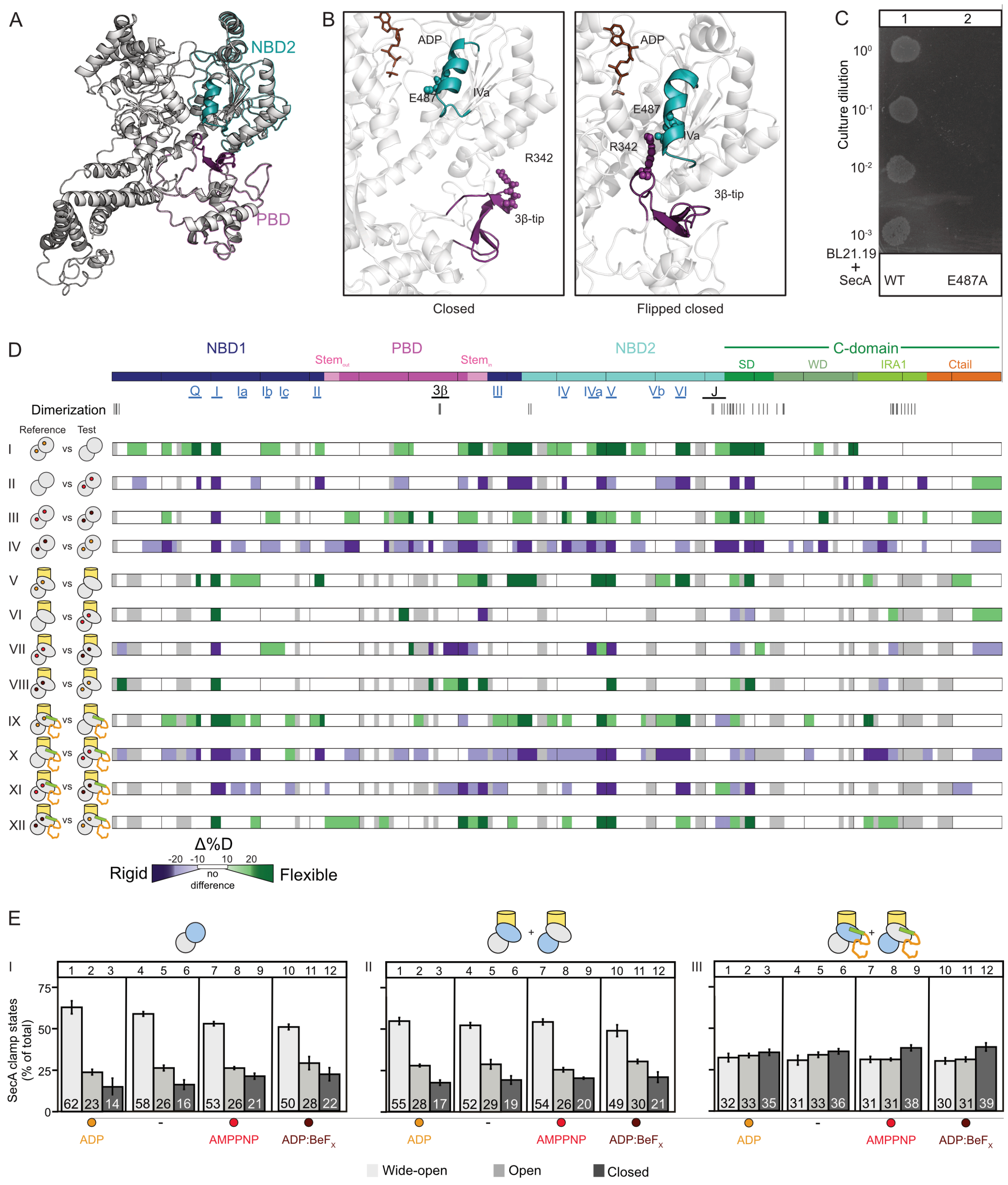


**Figure S6 Nucleotide regulated dynamics in channel-bound SecA and structure of the closed-flipped clamp state (related to Fig. 5)**

**A.** Structure of flipped-closed clamp state of SecA (PBD:3DIN) is shown with the 3β-tip (magenta) and motif IVa (cyan) highlighted. PBD (magenta) and NBD2 (cyan) are colour-outlined.

**B.** Close-up view of PBD-NBD interactions in the closed clamp state (as in Fig. S3, left) and flipped-closed clamp state of SecA (right; PBD:3DIN). The positions of $R342_{PBD}$ (3β-tip; magenta spheres) and $E487_{NBD2}$ (motif IVa; cyan spheres) are shown. R342 and E487 salt-bridge between them in the flipped-closed clamp state, making direct connections between the PBD and the helicase motifs in the ATPase motor.

**C.** In vivo genetic complementation assays of SecA and SecA(E487A) mutant, showing that E487 is essential for function. Experiments carried out as in (Fig. S4C).

**D.** Effect of nucleotide cycling on the local dynamics of SecA. ΔD-uptake differences between various nucleotide states are compared pairwise and presented as linear maps, as described (Krishnamurthy et al., 2021). Pairwise comparisons follow the dynamics of (I-IV) $SecA_2$ in solution, (V-VIII) SecYEG:$SecA_2$ and (IX-XII) SecYEG:$SecA_2$:preprotein as the translocase goes through the different nucleotide states of the ATP hydrolysis cycle. In each comparison, the reference state (left pictogram) is compared to the test state (right pictogram) with the differences in dynamics in the test state coloured (see scale; bottom). The domain organization of SecA is shown along with helicase motifs (in blue) and important clamp elements (in black). Residues involved in dimerization are indicated (Gouridis et al., 2013; Krishnamurthy et al., 2021).

Effect of nucleotides on $SecA_2$ in solution.

I. ADP release: $SecA_2$:ADP (reference) is compared to $SecA_2$ (test)

II. ATP binding: $SecA_2$ (reference) is compared to $SecA_2$:AMPPNP (test)

III. ATP transition state: $SecA_2$:AMPPNP (reference) is compared to $SecA_2$:ADP:$BeF_X$ (test)

IV. ATP hydrolysis: $SecA_2$:ADP:$BeF_X$ (reference) is compared to $SecA_2$:ADP

V-VIII. Effect of nucleotides on channel-primed $SecA_2$. Pairwise comparisons as in I-IV

IX-XII. Effect of nucleotides on channel-primed, preprotein activated $SecA_2$. Pairwise comparisons as in I-IV.

**E.** Effect of nucleotides on clamp dynamics of the translocase. Clamp states were quantified as described (Krishnamurthy et al., 2021) and shown (as in Fig. 3A). Clamp dynamics were determined for various translocase states in the presence of ADP (2 mM), *apo* state, AMPPNP bound (2 mM) and ADP:$BeF_X$ bound (1 mM). For channel-bound translocase states, the data is a composite of both active and regulatory protomers (Krishnamurthy et al., 2021).

(I) Clamp dynamics of solution $SecA_2$

(II) Clamp dynamics of channel-primed $SecA_2$

(III) Clamp dynamics of channel-primed preprotein-activated $SecA_2$

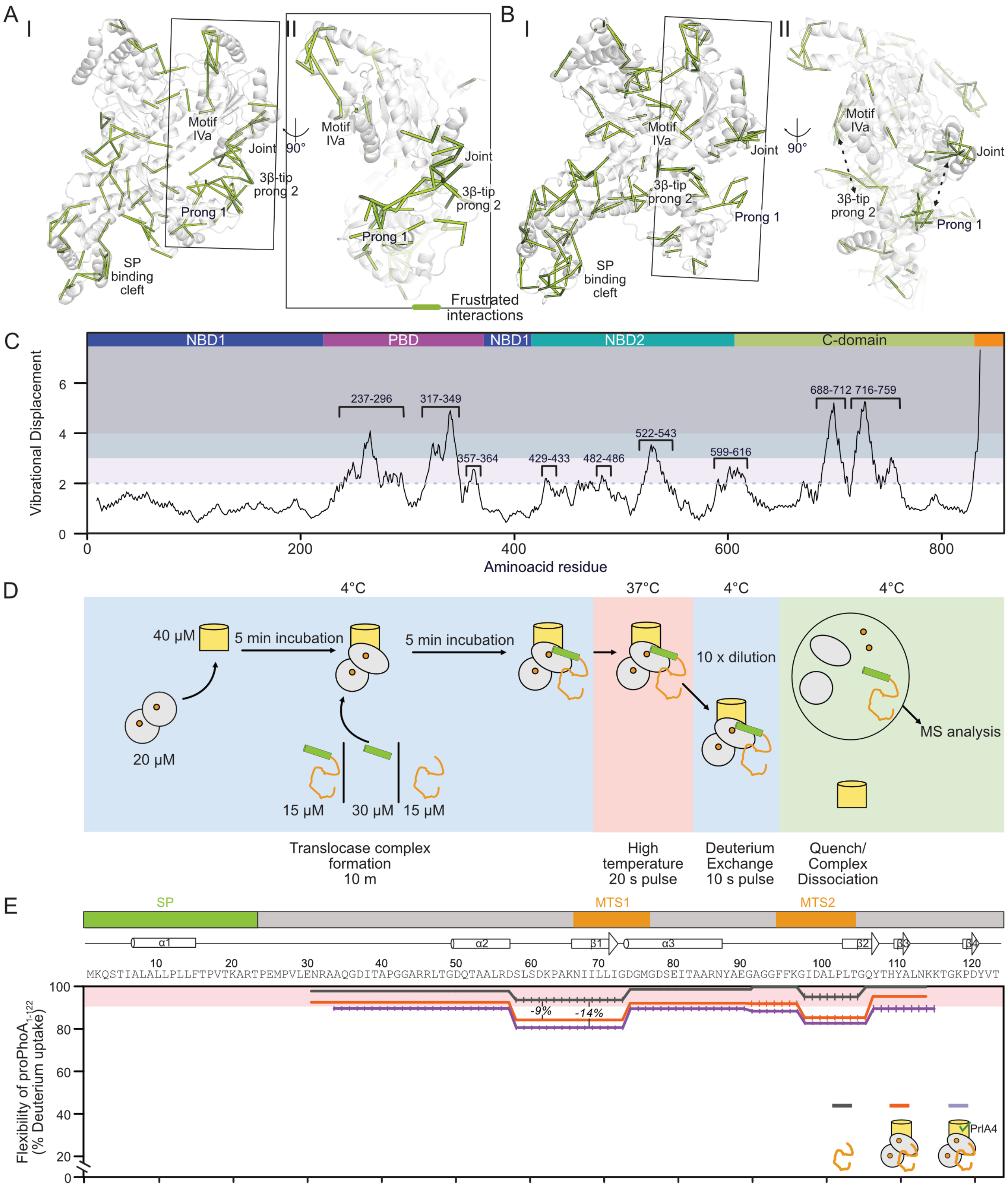


**Figure S7 Translocase binds and regulates preprotein dynamics (related to Fig. 6)**
**A-B.** Regions of frustration (green lines; as in Fig. 7B) derived from Frustratometer server (Parra et al., 2016) for the closed [**A**; structure derived from MD simulations (Krishnamurthy et al., 2021)] and closed flipped (**B**; structure modelled on PDB ID: 3DIN) structures. Structures are rotated 90°, focusing on frustrated regions in the clamp (enclosed in box) (**II**). Parallel lines of frustration are indicated (dashed black lines).

**C.** Per-residue vibrational displacement derived from normal mode analysis. Normal modes 7-12 (unweighted sum) were derived from the WebNM@ web server (Tiwari et al., 2014) using the PDB ID: 2VDA as the input *ec*SecA structure. Residues that undergo displacement greater than 2 (residue numbers provided) are highlighted in shades of blue (as in Fig. 6C) Per-residue displacement and normal mode analysis were carried out as described (Smit et al., 2021). Coloured SecA domains are indicated at the top of the graph.

**D.** Schematic representation of HDX-MS experiments following preprotein dynamics. Final concentration of reactants during complex formation are shown. The Sec translocase is assembled in the presence (or absence in *apo* conditions) of nucleotides with an excess of SecYEG. Client proteins are added in sub-stoichiometric amounts to the Sec translocase, to ensure all available client proteins are bound to the translocase. The activated complex is exposed to a 20 s pulse at 37 °C, and diluted (10 x dilution) in $D_2O$ buffer at 4 °C to start the D-exchange reaction. The D-exchange reaction is quenched with acid and the complex is dissociated in the presence of acidic proteases. The dynamics of the client proteins are analyzed by MS as described (Krishnamurthy et al., 2021).

**E.** Flexibility of $PhoA_{23\text{-}122}$ (as in Fig. 6A) as free (black line), bound to $SecYEG{:}SecA_2$ (orange line) or bound to $SecY_{PrlA4}EG{:}SecA_2$ (purple line).

#### Supplemental tables

##### Table S3 List of Buffers

| Buffer | Composition |
|---|---|
| **A** | 50 mM Tris-HCl, pH 8.0, 50 mM NaCl, 6 m Urea, 50% v/v glycerol |
| **B** | 50 mM Tris-HCl pH 8.0, 50 mM KCl, 1 mM $MgCl_2$, 1 mM DTT |
| **C** | 50 mM Tris-HCl pH 8.0, 50 mM KCl, 1 mM $MgCl_2$, 4 μM $ZnSO_4$, 2 mM TCEP |
| **D** | 1.3% formic acid, 4 mM TCEP, 1 mg/mL fungal protease XII |
| **E** | 50 mM MOPS pH 7.0, 50 mM KCl, 1 mM $MgCl_2$, 4 μM $ZnSO_4$, 2 mM TCEP |

##### Table S4 List of strains

| *E. coli* strain | Description (gene deleted) | Reference/source |
|---|---|---|
| DH5α | *F– Φ80lacZΔM15 Δ(lacZYA-argF) U169 recA1 endA1 hsdR17 (rK–, mK+) phoA supE44 λ– thi-1 gyrA96 relA1* | Invitrogen |
| BL21 (DE3) | T7 RNA polymerase gene under the control of the *lac* UV5 promoter | (Studier et al., 1990) |
| BL21.19 (DE3) | *secA13(Am) clpA::kan, ts* at 42°C; | (Mitchell and Oliver, 1993) |
| BL31 (DE3) | Non *ts*; spontaneous revertant of BL21.19 (DE3) | (Chatzi et al., 2017) |
| T7 Express lysY/I$^q$ Competent *E.coli* (High Efficiency) | Derivative of BL21 (DE3) deficient in Lon and OmpT proteases | New England Biolabs |

##### Table S5 List of plasmids

| Gene | Uniprot accession number | Plasmid name | Vector | Description/source/reference |
|---|---|---|---|---|
| *secA* | P10408 | pIMBB1280 | pET3a | (Gouridis et al., 2013) |
| His *secA 6-901* | P10408 | pIMBB7 | pET5a | (Gouridis et al., 2013) |
| - | | pLMB0081 | pET3a | pIMBB1280 was digested with NcoI, the *secA* N31-898 fragment was removed, the plasmid was re-ligated and used for *secA* NcoI fragment cloning |
| *secA*(Δα0/α1-6A) | P10408 | pIMBB1286 | pET3a | also called mSecA (Gouridis et al., 2013) |
| *secA*(V280C/L464) | P10408 | pLMB1646 | pET3a | also called SecA-D2 (Vandenberk et al., 2019) |
| His *secA* cys- | P10408 | pLMB1791 | pET16b | (Krishnamurthy et al., 2021) |
| His *secA* cys-(V280C/L464) | P10408 | pLMB1819 | pET16b | (Krishnamurthy et al., 2021) |
| His *secYEG* | P0AGA2 P0AG96 P0AG99 | pIMBB336 | pET610 | Gift from A. Driessen, University of Groningen, Groningen (van der Does, et al., 1998) |
| His *secY$_{PrlA4}$EG* | P0AGA2 P0AG96 P0AG99 | pIMBB842 | pET610 | (Gouridis et al., 2013) |
| *secA$_{PrlD23}$* | P10408 | pIMBB1314 | pET3a | also called PrlD23 |

| | | | | |
|---|---|---|---|---|
| | | | | PrI derivative of SecA with the Y134S mutation (Huie and Silhavy, 1995) introduced in secA 1-901 (Gouridis et al., 2013) |
| *secA*(H484Q) | P10408 | pLMB1858 | pET5a | Type I PrI derivative of SecA. *secA* N31-898 (H484Q) fragment from pIMBB578 (His SecA(H484Q) was inserted in pLMB0081 after NcoI digestion |
| His *secA*cys-(V280C/L464C/ H484Q) | P10408 | pLMB1907 | pET16b | The mutation H484Q was introduced in pLMB1819 using primer pairs X2154-X2155 |
| *secA*(H484A) | P10408 | pLMB1859 | pET5a | Type I PrI derivative of SecA. *secA* N31-898 (H484A) fragment from pIMBB527 (His SecA -H484A) was inserted in pLMB0081 after NcoI digestion |
| His *secA*cys-(V280C/L464C/ H484A) | P10408 | pLMB2027 | pET16b | The mutation H484A was introduced in pLMB1819 using primer pairs X930-X2238 |
| *secA*(L187A) | P10408 | pLMB1860 | pET5a | (Krishnamurthy et al., 2021) |
| His *secA*cys-(V280C/L464C/ L187A) | P10408 | pLMB1910 | pET16b | (Krishnamurthy et al., 2021) |
| His *secA*cys-(V280C/L464C/ A373V) | P10408 | pLMB2107 | pET16b | The mutation A373V was introduced in pLMB1819 using primer pairs X409-X911 |
| His *secA* N6-901 (L187A) | P10408 | pIMBB680 | pET5a | The mutation L187A was introduced in pIMBB7 using primer pairs X403-X901 |
| His *secA* N6-901 (L187V) | P10408 | pIMBB946 | pET5a | The mutation L187V was introduced in pIMBB7 using primer pairs X638-X639 |
| His *secA* N6-901 (L187I) | P10408 | pIMBB947 | pET5a | The mutation L187I was introduced in pIMBB7 using primer pairs X640-X641 |
| His *secA* N6-901 (A373V) | P10408 | pIMBB687 | pET5a | The mutation A373V was introduced in pIMBB7 using primer pairs X409-X911 |
| His *secA* N6-901 (A373I) | P10408 | pIMBB945 | pET5a | The mutation A373I was introduced in pIMBB7 using primer pairs X636-X637 |
| His *secA* N6-901 (A373F) | P10408 | pIMBB944 | pET5a | The mutation A373F was introduced in pIMBB7 using primer pairs X634-X635 |
| secA “LO” or His *secA* N6-834 (C98A/P301C/ S830C) | P10408 | pIMBB941 | pET5a | (Chatzi et al., 2017; Sardis et al., 2017) |
| secA “LC” or His *secA* N6-834 (K268C/I597C) | P10408 | pIMBB1394 | pET5a | (Chatzi et al., 2017; Sardis et al., 2017) |
| His *secA* N6-901 ($TGR_{342}AAA$) | P10408 | pIMBB701 | pET5a | The mutations T340A/G341A/R342A were introduced pIMBB07 using primer pairs X423-X1021 and X1768-X1769 |
| SecA E487A | P10408 | pLMB2110 | pET3a | The mutation E487A was introduced in pIMBB1280 using primer pairs X2373-X2374 |
| $proPhoA_{1-122}$ -His | P00634 | pIMBB1153 | pET22b | *prophoA* N1-122 fragment was isolated from pIMBB977 (*prophoA* Δcys His) using primers X560-X936 and inserted in pet22b after NdeI-XhoI digestion. |

| | | | | |
|---|---|---|---|---|
| $PhoA_{23-122}$-his | P00634 | pIMBB1183 | pET22b | *phoA* N23-122 fragment was isolated from pIMBB882 (*prophoA* His) using primers X806-X936 and inserted in pet22b after NdeI-XhoI digestion. |
| His *secA* N6-901 E181A | P10408 | pIMBB677 | pET5a | The mutation E181A was introduced in pIMBB7 using primer pairs X395-X400-X182 |
| His *secA* N6-901 F184A | P10408 | pIMBB678 | pET5a | The mutation F184A was introduced in pIMBB7 using primer pairs X395-X401-X182 |
| His *secA* N6-901 D185A | P10408 | pIMBB679 | pET5a | The mutation D185A was introduced in pIMBB7 using primer pairs X395-X402-X182 |
| His *secA* N6-901 R188A | P10408 | Pimbb681 | pET5a | The mutation R188A was introduced in pIMBB7 using primer pairs X395-X404-X182 |
| His *secA* N6-901 D189A | P10408 | pIMBB682 | pET5a | The mutation D189A was introduced in pIMBB7 using primer pairs X395-X405-X182 |
| His *secA* N6-901 M191A | P10408 | pIMBB683 | pET5a | The mutation M191A was introduced in pIMBB7 using primer pairs X395-X406-X182 |
| His *secA* N6-901 M191F | P10408 | pIMBB948 | pET5a | The mutation M191F was introduced in pIMBB7 using primer pairs X642-X643 |

#### Table S6 List of primers

| # | Description | **DNA sequence** (5'-3'; Mutated codons in bold; restriction sites underlined) |
|---|---|---|
| **X182** | Reverse *secA* primer for mega-primer mutagenesis | GGCCTTTCGCAGTACGTTC |
| **X395** | Forward primer to amplify from position 4190 to 4214 of pIMBB7 for mega-primer mutagenesis | CTAACAACAATAAACCTTTACTTC |
| **X400** | Reverse mutagenic primer for SecA E181A | GTCAAAGCCGTA**TGC**GTTGTTCGT |
| **X401** | Reverse mutagenic primer to generate *secA* F184A | GCGCAGGTAGTC**GGC**GCCGTATTCGTTGTT |
| **X402** | Reverse mutagenic primer to generate *secA* D185A | GTCGCGCAGGTA**GGC**AAAGCCGTATTCGTTGTT |
| **X403** | Reverse mutagenic primer to generate *secA* L187A | CATGTTGTCGCG**CGC**GTAGTCAAAGCCGTA |
| **X404** | Reverse mutagenic primer to generate *secA* R188A | CATGTTGTC**GGC**CAGGTAGTCAAA |
| **X405** | Reverse mutagenic primer to generate *secA* D189A | GAACGCCATGTT**CGC**GCGCAGGTAGTCAAA |
| **X406** | Reverse mutagenic primer to generate *secA* M191A | CAGGGCTGAACGC**CGC**GTTGTCGCG |
| **X409** | Forward mutagenic primer to generate *secA* A373V | GAAAACCAAACGCTG**GTT**TCGATCAC |
| **X423** | Forward mutagenic primer generate *secA* T340A/G341A/R342A | GACGAACAC**GCCGCTGCT**ACCATGCAGGG |
| **X560** | Forward phoA primer inserting an NdeI restriction site | GGGAATTCCATATAAACAAAGCACTATTGCA |
| **X634** | Forward mutagenic primer to generate *secA* A373F | GAAAACCAAACGCTG**TTT**TCGATCACCTTCCAG |

| | | |
|---|---|---|
| **X635** | Reverse mutagenic primer to generate *secA* A373F | CTGGAAGGTGATCGAAAACAGCGTTTGGTTTTC |
| **X636** | Forward mutagenic primer to generate *secA* A373I | GAAAACCAAACGCTGATTTCGATCACCTTCCAG |
| **X637** | Reverse mutagenic primer to generate *secA* A373I | CTGGAAGGTGATCGAAATCAGCGTTTGGTTTTC |
| **X638** | Forward mutagenic primer to generate *secA* L187V | TACGGCTTTGACTACGTGCGCGACAACATGGCG |
| **X639** | Reverse mutagenic primer to generate *secA* L187V | CGCCATGTTGTCGCGCACGTAGTCAAAGCCGTA |
| **X640** | Forward mutagenic primer to generate *secA* L187I | TACGGCTTTGACTACATCCGCGACAACATGGCG |
| **X641** | Reverse mutagenic primer to generate *secA* L187I | CGCCATGTTGTCGCGGATGTAGTCAAAGCCGTA |
| **X642** | Forward mutagenic primer to generate *secA* M191F | TACCTGCGCGACAACTTCGCGTTCAGCCCTGAA |
| **X643** | Reverse mutagenic primer to generate *secA* of M191F | TTCAGGGCTGAACGCGAAGTTGTCGCGCAGGTA |
| **X806** | Forward phoA primer annealing at Thr 23, inserting NdeI and HindIII restriction sites | GGGAATTCCATATGAAGCTTACACCAGAAATGCCTGTTCTGGAA |
| **X901** | Forward mutagenic primer to generate *secA* L187A | TACGGCTTTGACTAC**GCG**CGCGACAACATG |
| **X911** | Reverse mutagenic primer to generate *secA* A373V | GTGATCGA**AAC**CAGCGTTTGGTTTTC |
| **X930** | Forward mutagenic primer to generate *secA* H484A | TGAACAACGCCAAATTC**GCC**GCCAACGAAGCG |
| **X936** | Reverse phoA primer annealing at Thr 122, inserting a XhoI site, a Tyrosine and a stop codon | GACCCGCTCGAG**TTAATA**GGTGACGTAGTCCGGTTTG |
| **X1021** | Reverse mutagenic primer to generate *secA* T340A/G341A/R342A | CCCTGCATGGT**AGCAGCGGC**GTGTTCGTC |
| **X1768** | Forward mutagenic primer to generate *secA* L464C | CCATCGAAAAATCGGAGTGCGTGTCAAACGAACTG |
| **X1769** | Reverse mutagenic primer to generate *secA* L464C | CAGTTCGTTTGACACGCACTCCGATTTTTCGATGG |
| **X2373** | Forward mutagenic primer to generate *secA* E487A | AAATTCCACGCCAACGCCGCGGCGATTGTTGCT |
| **X2374** | Reverse mutagenic primer to generate *secA* E487A | AGCAACAATCGCCGCCGGGTTGGCGTGGAATTT |

**Supplementary Movies**

**Movie S1** Clamp motions of the translocase (related to Fig. 3 and S6)

Movie depicting clamp motion from Wide-open state to closed flipped state. Movies were generated using PyMOL using *E.coli* homology models (Sardis and Economou, 2010) of wide open (PDB: 1M6N), open (PDB:2VDA) and closed flipped PBD (PDB:3DIN) states.
